# GIV•Kindlin interaction is required for Kindlin-Mediated Integrin Recognition and Activation

**DOI:** 10.1101/870113

**Authors:** Cristina Rohena, Nicholas Kalogriopoulos, Navin Rajapakse, Suchismita Roy, Inmaculada Lopez-Sanchez, Jailal Ablack, Debashis Sahoo, Pradipta Ghosh

## Abstract

Cells perceive and respond to the extracellular matrix (ECM) *via* integrin receptors; their dysregulation has been implicated in inflammation and cancer metastasis. Here we show that a guanine nucleotide exchange modulator of trimeric-GTPase Gαi, GIV (*a*.*k*.*a* Girdin), directly binds the integrin adaptor Kindlin-2. A non-canonical short linear motif within GIV’s C-terminus binds Kindlin-2-FERM3 domain at a site that is distinct from the binding site for the canonical NPxY motif on the -integrin tail. Binding of GIV to Kindlin-2 allosterically enhances Kindlin-2’s affinity for β1-integrin. Consequently, integrin activation and clustering are maximized, which augments cell adhesion, spreading and invasion. Findings elucidate how the GIV•Kindlin-2 complex has a two-fold impact: it allosterically synergizes integrin activation and enables β1-integrins to indirectly access and modulate trimeric GTPases *via* the complex. Furthermore, Cox proportional-hazard models on tumor transcriptomics provide trans-scale evidence of synergistic interactions between GIV•Kindlin-2•β1-integrin on time to progression to metastasis.

**The eTOC blurb:** Integrins mediate cell adhesion to the extracellular matrix; their dysregulation fuels inflammation, cancer cell invasion and metastasis. Authors show how two pro-metastatic scaffold proteins, Kindlin and GIV/Girdin bind and cooperatively enhance their allosteric coupling to integrins, and their subsequent activation. Findings reveal novel interfaces in integrin signaling for pharmacologic manipulation.

**HIGHLIGHTS:**

- GIV and Kindlin(K2), two integrin adaptors that promote metastasis, bind each other
- Binding of GIV or integrin to K2 allosterically enhances GIV•K2•integrin complexes
- Binding is required for the maximal recruitment of GIV and K2 to active integrins
- Binding facilitates integrin clustering, activation, tumor cell adhesion, invasion.

## INTRODUCTION

Cell adhesion to extracellular matrix (ECM) proteins and to neighboring cells is essential for multicellular organisms. Cells perceive and respond to the composition and stiffness of the ECM through mechanotransduction *via* integrin class of receptors. As key regulators of cell fate, e.g., survival, proliferation and migration, integrins are essential for embryonic development and their aberrant signaling fuels numerous diseases, including inflammation, tumor growth, chemoresistance and metastasis.

Several adaptor proteins transduce integrin signals by directly binding to the cytoplasmic tails of β-integrins (Legate and Fassler, 2009). Among them, only talin and kindlin are known to be indispensable for integrin activation (Calderwood et al., 2013; Theodosiou et al., 2016). The phosphotyrosine-binding (PTB)-like module in talin’s FERM3 domain binds to the membrane-proximal NPxY motif of β-integrin cytoplasmic tails. Kindlin is also a FERM3 domain-containing protein that binds to the membrane distal NPxY motif on integrin tails. Structural, biochemical and mutational approaches revealed that talin-mediated integrin activation results in conformational changes in the receptor’s transmembrane helix (Lau et al., 2008a; Lau et al., 2008b), and how such activation requires first the reversal of an intramolecular autoinhibitory contact within talin-- between its FERM3-PTB and its rod domains (Garcia-Alvarez et al., 2003; Goksoy et al., 2008; Vinogradova et al., 2002; Wegener et al., 2007). By contrast, kindlin does not have the ability to directly alter the conformation of the integrin transmembrane helix and instead augments integrin activation in cooperativity with talin (Bledzka et al., 2012), without which the conformational shift of integrins from the low-to high-affinity state cannot occur and maximal activation is not possible (Calderwood et al., 2013; Campbell and Humphries, 2011)].

Regarding mechanistic insights into how adaptor binding translates to integrin activation, gathering evidence suggests that activation occurs by finetuning the affinity of talin and kindlin for integrins. Talin’s affinity for integrins, for example, can be regulated either through proteolytic relief of auto-inhibition or through membrane recruitment and molecular allostery upon direct binding of phosphoinositides to the FERM3-domain (Han et al., 2006; Lagarrigue et al., 2016; Lee et al., 2009; Moore et al., 2012; Ye et al., 2016). In the case of kindlin, however, despite the discovery of numerous interacting partners (Bottcher et al., 2017; Fukuda et al., 2014; Theodosiou et al., 2016) that coordinate the activation of focal adhesion kinase (FAK), Rac1 and the Arp2/3 complex within the so-called adhesome complex (Bottcher et al., 2017; Theodosiou et al., 2016; Winograd-Katz et al., 2014; Zaidel-Bar et al., 2007), who or what may finetune the affinity of kindlin for integrins remains unknown and whether such finetuning enhances initial integrin activation, clustering and adhesion strengthening remains elusive.

Besides talin and kindlin, we (Lopez-Sanchez et al., 2015) and others (Leyme et al., 2016; Leyme et al., 2015) have recently reported that exposing cells to ECM also triggers the tyrosine phosphorylation of another adaptor protein, GIV and that focal adhesions (FAs) serve as the major hubs for tyr-based mechanochemical signaling via GIV. GIV is a guanine nucleotide exchange factor (GEF) for trimeric GTPase, Gi (Garcia-Marcos et al., 2009; Kalogriopoulos et al., 2019) and is an actin remodeler (Enomoto et al., 2005). Published work has shown that GIV (and Gαi via GIV) interact exclusively with ligand-activated β1-integrins and GIV’s GEF function is essential for the subsequent activation of Gαi in response to stimuli (Leyme et al., 2015; Lopez-Sanchez et al., 2015). GIV-dependent Gi activation and release of ‘free’ Gβγ-heterodimer modulates multiple downstream signals including FAK activation, remodeling of the actin cytoskeleton, and Rac1 and PI3K-dependent signaling, resulting in enhanced haptotaxis and invasion (Leyme et al., 2015). These signals and cellular phenotypes are further reinforced *via* a forward-feedback loop in which activated FAK phosphorylates GIV and that such phosphorylation further enhances PI3K-Akt signaling, the integrity of FAs, cell adhesion and motility (Leyme et al., 2016; Lopez-Sanchez et al., 2015). Spatially restricted signaling via tyrosine phosphorylated GIV at the FAs is enhanced during cancer metastasis (Midde et al., 2018). Despite these insights, how GIV binds integrins remained unclear; because they co-immunoprecipitated from cells, but recombinant GIV and β-integrin cytoplasmic tails did not interact *in vitro*, the interaction was assumed to be indirect (Lopez-Sanchez et al., 2015).

We set out to investigate how GIV gains access to β-integrin cytoplasmic tails, but unexpectedly stumbled upon an intermolecular interplay between GIV, kindlin and β1-integrin. Findings help answer some of the fundamental unanswered questions e.g., how GIV may bind integrins to impact downstream signaling and how kindlin may bring about maximal integrin activation; they also provide mechanistic insights into how cooperativity between the two adaptors may be essential for both.

## RESULTS AND DISCUSSION

### GIV interacts with ligand-activated β1-integrins indirectly via kindlin-2

Prior work had shown that GIV indirectly associates with β1-integrins early (within minutes) during cell adhesion and localizes to nascent focal adhesions (FAs) at the cell periphery well before integrins cluster and FAs mature. Because integrin activation and clustering, two of the earliest steps of cell adhesion, require sequential recruitment of the adaptor proteins talin, then kindlin, and finally, tensin (Bachir et al., 2014; Calderwood et al., 2013; Montanez et al., 2008), we hypothesized that GIV may interact with one or more of these adaptors. To test this hypothesis, we carried out pulldown studies using recombinant GST-tagged integrin-binding FEM3/PTB modules of the key adaptor proteins (Talin, Kindlin and Tensin) and His-tagged GIV’s C-terminal fragment (GIV-CT; ∼ 210 aa). To avoid common problems encountered when generating protein fragments (misfolding, degradation, unforeseen/undesired mutations, all leading to non-functional proteins, we refrained from creating new constructs and instead curated previously published constructs that were validated in various biochemical or crystallography studies to interrogate integrin biology and sequenced them to confirm accuracy (see *STAR methods*). As for GIV-CT, we have previously been shown to bind cytoplasmic tails of multiple growth factor receptors (Lin et al., 2014), G proteins (Garcia-Marcos et al., 2009), FAK (Lopez-Sanchez et al., 2015) and p85α-PI3K (Lin et al., 2011); prior work also showed that GIV-CT is sufficient to trigger cancer cell invasion (Ma et al., 2015a; Midde et al., 2015). Among the three Kindlins (K1– 3), we chose to study kindlin-2 (K2) because unlike K1 and K3, which are expressed in restricted cells/tissues (i.e., epithelial and hematopoietic system, which express GIV at very low levels; (Enomoto et al., 2005)), K2 is expressed ubiquitously (Rognoni et al., 2016). We found that GIV specifically bound the FERM3-PTB module of K2 and to the PTB module of tensin, but not to the FERM2-PTB module of talin [**Fig 1A**]. Because GIV was previously shown to accumulate early during cell adhesion within nascent FAs at the cell periphery (Lopez-Sanchez et al., 2015), where K2 is known to exist at a 1:1 ratio with β1 integrins (Sun et al., 2014), and tensin on the other hand is typically enriched later in FAs during the course of tension-dependent maturation (Zaidel-Bar et al., 2004), we chose to further dissect the nature and relevance of the GIV•K2 interaction. Full length GIV (from cell lysates) could also bind GST-K2, but little or no binding was observed with GST-β integrin tails [**Fig 1B**]. We also confirmed that the reverse was also true, in that, when GST-GIV-CT was immobilized on glutathione beads, it could directly bind His-K2 [**Fig 1C; S1A**]; to our surprise, in this assay, K2 bound GIV-CT much better than it bound our positive control (GST-β1-CT). Because one prior study (Leyme et al., 2015) claimed the existence of a possible weak, but direct interaction between GIV’s N-terminus (GIV-NT; aa 1-256) and the cytoplasmic tail of β1-integrin, we compared side by side the ability of the N- and C-terminal fragments to bind GST-β1-CT and K2. We found that neither the C-, nor not the N-terminus of GIV bound the cytoplasmic tail of integrin β1 to any appreciable degree when comparable with the binding of GIV-CT to K2 [**Fig S1B**]. Furthermore, co-immunoprecipitation studies confirmed that the interaction we observe between tagged recombinant proteins *in vitro* occurs also between full-length endogenous GIV and kindlin2 proteins in cells [**Fig 1D**]. These findings demonstrate that GIV’s C-terminus directly and specifically binds the integrin-adaptor K2 and suggest that previously reported interactions of GIV and G protein, Gi with ligand-activated β1-integrin in cells are likely to be indirect (*via* K2). Indeed, we confirmed this to be true, because neither GIV, nor Gαi was detectable within β1-integrin-bound complexes when we depleted HeLa cells of endogenous K2, but they were readily detectable when an siRNA-resistant GFP-tagged K2 was exogenously expressed in these cells [**Fig 1E**]. While it is possible that impaired formation of focal adhesions in cells without kindlin-2 may preclude the GIV•β1 integrin interaction, together with the biochemical evidence of direct interaction and interactions observed in cells, our findings indicate that GIV interacts with ligand-activated β1-integrins indirectly via kindlin-2.

**Figure 1.**
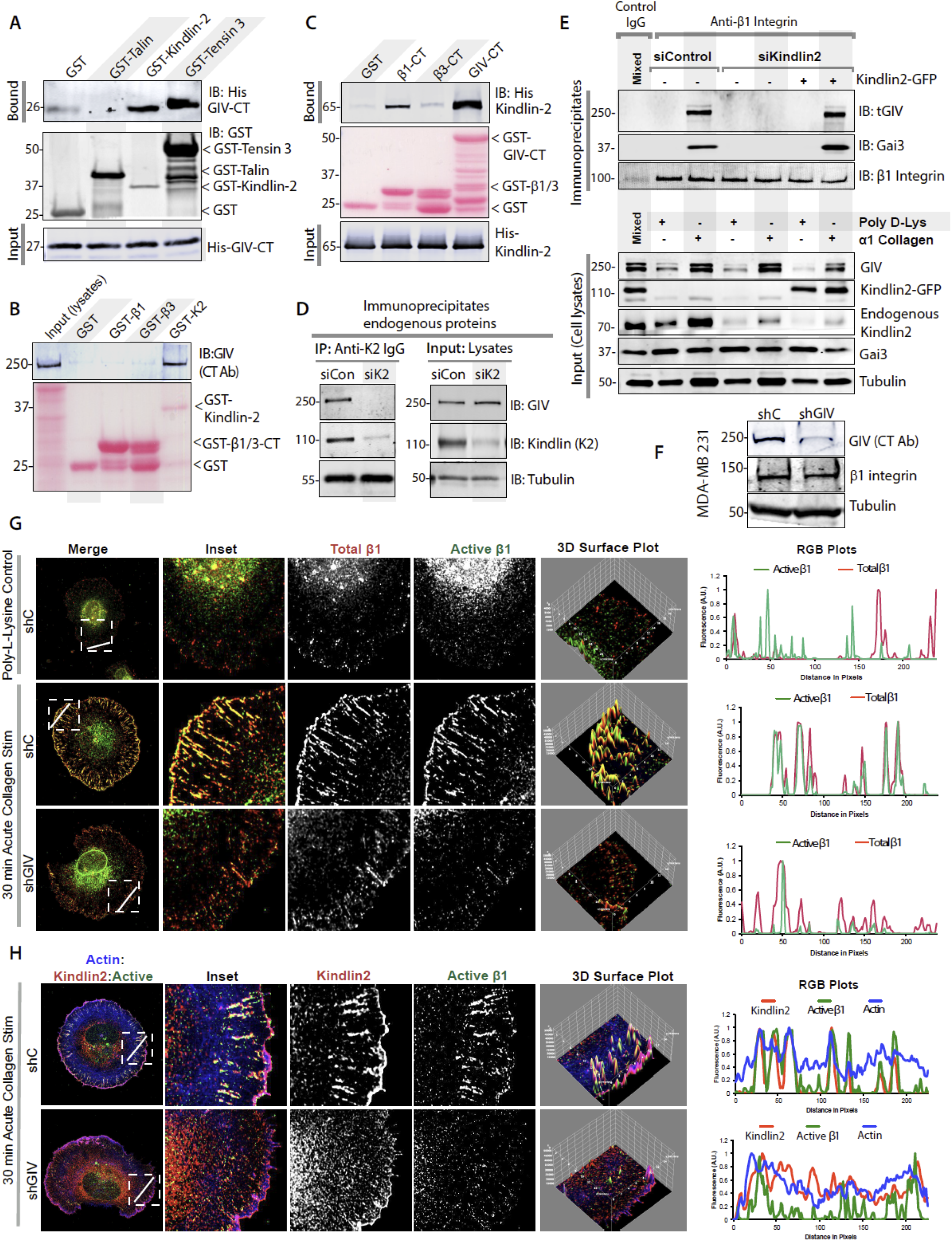
GIV binds kindlin-2, is required for kindlin recruitment to integrin β1 at focal adhesions, and for maximal integrin activation. **(A)** GST pulldown assays were performed using equal aliquots of recombinant His-GIV-CT (aa 1660-1870; ∼3 μg) and various FERM3/PTB fragments of GST-tagged adaptor proteins known to bind integrin-β receptors. Bound His-GIV-CT and various GST ligands were visualized by immunoblotting using an anti-His mAb and an anti-GST pAb, respectively. **(B)** Equal aliquots of lysates of MDA-MB-231 cells were used as source for full-length endogenous GIV in GST pulldown assays with GST-Kindlin 2 (K2), GST-β1 or GST-β3 proteins. Bound GIV was visualized by immunoblotting using anti-GIV-CT rabbit polyAb. Equal loading of GST proteins were confirmed by Ponceau S staining. **(C)** GST pulldown assays were performed using recombinant His-Kindlin 2 [6xHis-SUMO-Kindlin2Δ (Li et al., 2017)] and GST-GIV-CT (aa 1623-1870). Bound kindlin-2 was analyzed by immunoblotting with anti-His mAb. GST proteins were visualized by Ponceau S staining. GST-β1 and GST-β3 integrin tails were used as positive and negative controls, respectively. See also **Fig S1A** for bar graphs displaying quantification. **(D)** Immunoprecipitation studies with anti-Kindlin2 (K2) antibody on lysates of control (siC) or kindlin-2-depleted (by siKindlin-2) HeLa cells. Immune complexes were analyzed for GIV and Kindlin2 by immunoblotting (IB). **(E)** Control (siC) or kindlin-2-depleted (by siKindlin-2) HeLa cells reconstituted or not with GFP-Kindlin-2, grown on poly-D-Lysine-coated surface were stimulated (+) or not (–) by plating on collagen-coated surface for 30 min before lysis. Equal aliquots of lysates (Input cell lysates) were subjected to immunoprecipitation using anti–β1 integrin antibody. Immune complexes were analyzed for total (t)GIV, β1 integrin, and Gαi3 by immunoblotting. The differences in levels of tGIV in input lysates likely reflects differential extraction by Triton-X100 in cells plated on poly-D-Lysine vs collagen-plated coverslips. **(F)** MDA-MB-231 cells depleted (shGIV) or not (shC) of GIV by shRNA were analyzed for GIV and tubulin by Immunoblotting with respective antibodies. **(G-H)** Cell lines in F were grown on poly-D-Lysine-coated surface were stimulated by plating on collagen-coated cover slips for 30 min, fixed and stained for active β1 integrin (green; using a conformational specific antibody, clone 9EG), phalloidin (blue; actin), and either total β1-integrin (G; red) or K2 (H; red) and analyzed by confocal microscopy. Representative deconvolved images are shown. Insets were analyzed by rendering 3D surface plots and line scans were taken to generate RGB plots using ImageJ.

### GIV is required for integrin activation, formation of β1-integrin•Kindlin complexes at focal adhesions

Because K2 is believed to be indispensable for integrin activation (Calderwood et al., 2013; Theodosiou et al., 2016), next we analyzed if GIV, which is co-recruited with K2 is also required for the same. To this end, we used control and GIV-depleted the MDA-MB-231 breast cancer cells, a model system that was previously used by others to implicate GIV’s role in integrin-dependent downstream activation of the PI3K-Akt and RhoA pathways (Leyme et al., 2015), stimulated them by plating on collagen-coated cover slips and stained them with a previously validated conformation-sensitive rat anti-CD29 9EG7 antibody. We found that cells without GIV had significantly reduced integrin activation, both within nascent FAs in the cell periphery and within mature FAs [**Fig 1G-H**], indicating that much like K2, GIV is also required for β1-integrin activation. When we monitored by immunofluorescence the recruitment of endogenous K2 in these cells, we unexpectedly found that the colocalization of active-β1 and K2 was also reduced in GIV-depleted cells [**Fig 1H**], indicating that GIV may somehow enable the formation of active β1-integrin•K2 complexes at the FAs. Although the levels of total β1-integrin were reduced at the cell periphery in GIV-depleted cells, immunoblots revealed that the levels of the protein was not reduced (**Fig 1F**), instead redistributed to vesicular structures at the center of the cell, which could represent some endocytic compartment. Although GIV-depleted cells did not always spread as well (quantified later in-depth), the findings in **Fig 1G-H** were observed in GIV-depleted cells that had spread to an equivalent degree as control cells, indicating that GIV may impact integrin activation and K2 recruitment to integrins regardless of cell spreading. Taken together, we conclude that GIV is recruited to the cytoplasmic tails of β1-integrin indirectly *via* its ability to bind the K2 adaptor, and that GIV may be a necessary component of integrin activation and for the formation of active-β1•K2 complexes.

### GIV’s C-terminus binds kindlin’s FERM3-PTB domain via a non-canonical short linear motif (SLIM)

β1-integrins bind the FERM3-PTB modules of talin and kindlin *via* canonical mechanisms that involve two conserved NPxY motifs on the cytoplasmic tail of the receptor. Because GIV lacks a similar motif, we hypothesized that the mechanism of GIV•K2 interaction may be non-canonical. The first clues into the mechanism came from a previously solved NMR structure of tensin-PTB) bound to a peptide derived from the protein, Deleted in Liver Cancer (DLC1) revealing a non-canonical mode of binding to PTB/FERM3-PTB modules (Chen et al., 2012). A sequence alignment showed that the core sequence “Pro(P)-Gly(G)-x-Phe(F)” in DLC1 that was previously implicated in binding tensin-PTB was conserved within GIV’s C-terminus [**Fig 2A**]; the Phe(F) within this motif was determined to contribute significantly to the strength of the interaction by filling a shallow pocket within the PTB domain of tensin. This putative PTB-binding short linear interaction motif (SLIM) in GIV-CT is situated downstream of the GEM motif that GIV uses to bind and activate Gαi and is within a stretch of sequence that was predicted to have the highest degree of disorder [**Fig S2A-C**]. This region has previously been shown to also fold into an SH2-like module upon recognizing phosphotyrosines on the cytoplasmic tails of diverse ligand-activated growth factor RTKs (Lin et al., 2014) [**Fig S2D**]. This motif also appeared to be evolutionarily young, i.e., conserved only in higher mammals [**Fig 1C**], much like GIV’s SH2-like module (Lin et al., 2014).

We asked if the putative PTB-binding SLIM in GIV-CT is functional. First, using GST-tagged SH2 and PTB fragments of tensin, we confirmed that full length GIV specifically binds the PTB domain of tensin [**Fig 2D**]. We found that the WT, but not a mutant His-GIV-CT protein in which the Phe(F) within the PGxF sequence is mutated to alanine (PGxA) could bind GST-Tensin-PTB [**Fig 2E**], which confirmed that the tensin-PTB•GIV-CT interaction is direct that it requires the intact PGxF SLIM. When we carried out similar assays, but replaced GST-tensin-PTB with GST-K2-FERM3-PTB, we observed identical results [**Fig 2F-G**], which confirmed that the K2-FERM3-PTB•GIV-CT interaction is also direct, and that it too requires the intact PGxF SLIM. Using pulldown assays with GST-K2 and an array of His-tagged GIV-CT mutants that perturb the core PGxF SLIM, we further determined the relative contributions of the various residues within the SLIM: mutating the Pro(P) and Gly(G) to Ala(A) had a partial effect on the interaction, while mutating the Phe(F) was the most disruptive [**Fig 2H**]. As expected, based on the location of the PGxF motif within GIV’s C-terminus [**Fig S2D**], none of the mutants impacted GIV’s ability to bind Gαi [**Fig S3**]. These studies provided the rationale for the use of the PGxA single point mutant in all further studies as a precise tool to specifically dissect the functional relevance of the GIV•K2 interaction.

**Figure 2.**
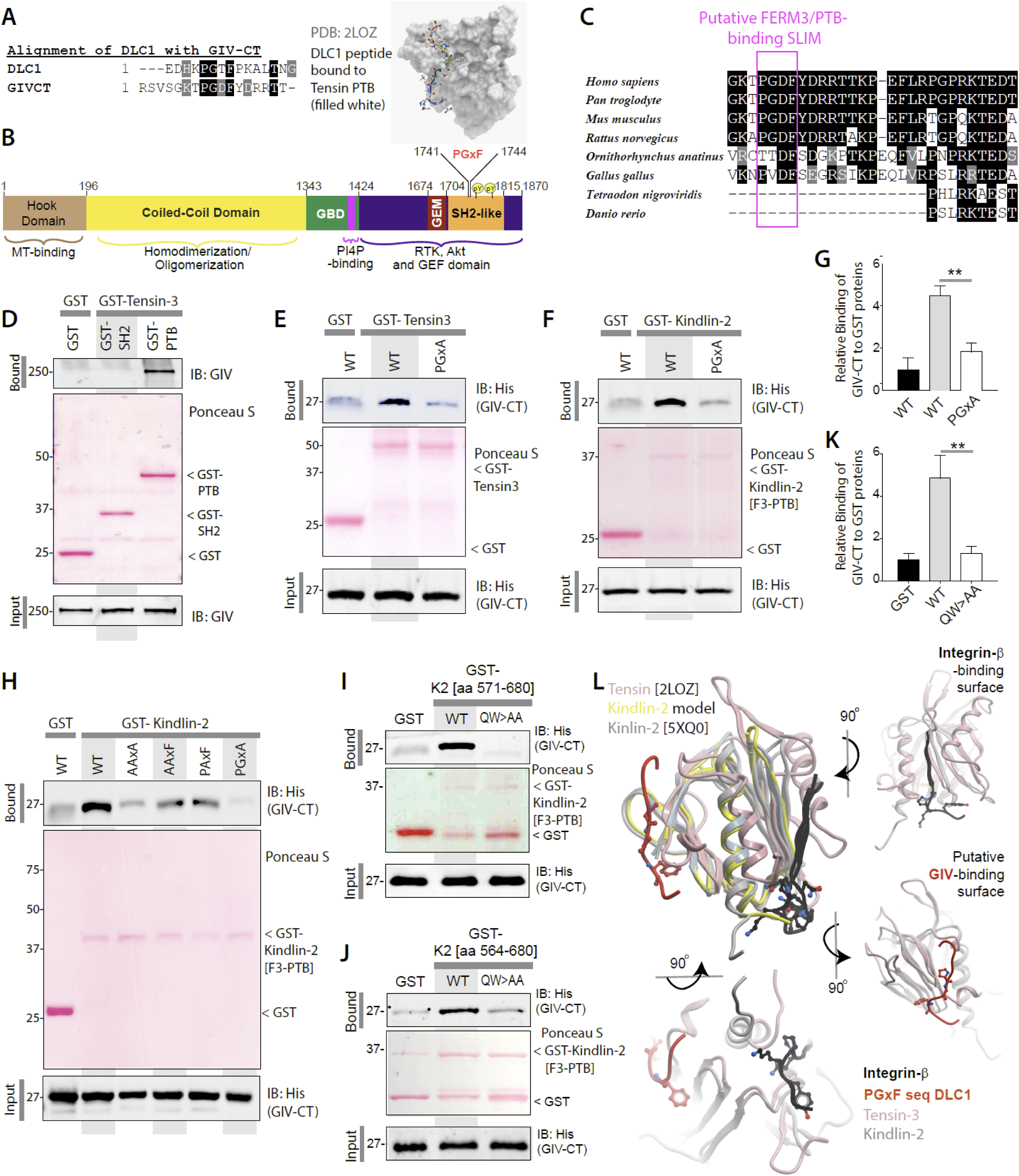
Identification and validation of a short linear motif (SLIM) in GIV that directly binds the FERM3(F3)-PTB module in Kindlin-2 (K2). **(A)** Sequence alignment between GIV’s C-terminus and DLC1 (which binds Tensin *via* non-canonical mechanisms) using ClustalW and Boxshade revealed a conserved putative SLIM [PGxF] (Left) which was implicated in binding the PTB module of Tensin (NMR of DLC1-bound Tensin co-complex; Right). **(B)** Bar diagram showing the various modules in GIV. The PGxF SLIM is located within the C-terminal stretch of GIV, which is largely disordered (see **Figure S2A-C**). **(C)** Sequence alignment of GIV’s C-terminus shows that the PGxF SLIM is evolutionarily young (only conserved in higher organisms). The relative positions of this and other SLIMs (that bind G proteins, PI3-kinase) and modules (that facilitate RTK binding) within the intrinsically disordered C-terminus are shown in **Figure S2D**. **(D)** GST pulldown assays were performed using lysates of Cos7 cells as source of full length GIV with GST-Tensin-3 SH2 and SH2+PTB fragments. Bound proteins were analyzed by immunoblotting with anti-His mAb. **(E)** GST pulldown assays were performed using WT or ^1741^PGxA^1744^ [F1744A] mutant His-GIV-CT and GST-Tensin-3 (SH2+PTB). **(F)** GST pulldown assays were performed using WT or ^1741^PGxA^1744^ [F1744A] mutant His-GIV-CT and GST-Kindlin-2 (F3-PTB). **(G)** Bar graphs display the relative binding of His-GIV-CT to GST proteins in F. Error bar = S.E.M (n = 5); **p=<0.01. **(H)** GST pulldown assays were performed using of His-GIV-CT WT or various mutants targeting its PGxF sequence and GST-Kindlin-2 (F3-PTB). **(I-K)** GST pulldown assays were performed using GST-K2 [aa 571-680, I and aa 564-680, J] WT or a QW/AA mutant proteins immobilized on glutathione beads and His-GIV-CT (aa 1660-1870). Bound proteins were analyzed as in D. Bar graphs (K) display the relative binding of His-GIV-CT to GST-K2 proteins in H and I. Error bar = S.E.M (n = 5); **p=<0.01. **(L)** A structural model built using the solved structures of DLC1-bound Tensin and Integrin-bound K2 as templates predicted that the FERM3-PTB module of K2 (grey ribbon) may simultaneously bind the non-canonical PGxF sequence on GIV’s C-terminus (red) as well as the canonical NPxY sequence on the cytoplasmic tail of β1-integrins (black) on via two distinct binding surfaces. These surfaces are exposed also when K2 is dimerized (see **Fig S4**).

We then asked if perturbing the PTB-like conformation within the FERM3 module of K2 impacts the GIV•K2 interaction. To this end, we used a previously validated mutant in which Glu(Q)^614^Try(W)^615^ is mutated to AA in the FERM3 subdomain of K2, which impairs PTB-like folding and recognition of canonical NPxY sequences on β-integrins (Ma et al., 2008; Shi et al., 2007). Defective integrin activation in cells expressing this K2 mutant has previously been attributed to its inability to assemble the K2•β-integrin interface (Bledzka et al., 2016; Xu et al., 2014) (Ma et al., 2008; Montanez et al., 2008; Shi et al., 2007). Using two different GST-K2 constructs varying slightly in their construct boundaries [one K2 construct (aa 571-680)was used by others to characterize the K2•β1 interaction *in vitro* and in cells (Montanez et al., 2008) and another construct was generated by us (aa 568-680) and spans the complete FERM3 module], we found that this PTB-defective QW>AA mutant K2 is required for not just binding the NPxY sequence on β-integrins, but also for binding GIV, indicating that the QW>AA K2 mutant is non-selective, in that, the mutations impair both interactions indiscriminately. Because multiple groups observed impaired integrin activation in cells expressing the QW>AA K2 mutant, and we observe similar defects in activation in GIV-depleted cells [**Fig 1G-H**], it is possible that the observed defect in the QW>AA K2-expressing cells is not just due to impaired K2•β-integrin interaction, but equally likely to be due to impaired K2•GIV interaction. Hence, to specifically study the impact of the GIV•K2 interaction, we used the newly identified PGxA point mutant of GIV that is deficient in binding to K2 in all subsequent assays.

### The GIV•kindlin-2 interaction allosterically enhances kindlin’s affinity for β1-integrin

Next we asked if and how binding of GIV to K2 impacts the K2•β1-integrin interaction. Three possible scenarios were considered: 1) GIV and β1-integrin may compete for the same site or bind on overlapping sites on K2, and if so, their interactions will be mutually exclusive; 2) they may each bind K2 non-competitively *via* two interfaces without steric hindrances, and if so, they may bind concurrently and exist as ternary GIV•K2•β1-integrin complexes; 3) they may bind non-competitively at distinct sites on K2, but allosterically impact (either inhibit or augment) each other’s’ ability to bind K2. Superimposition of the DLC1(PGxF)•tensin complex structure (Chen et al., 2012), the recently solved β1•K2 co-complex structure (Li et al., 2017) and a homology model of K2 (built using tensin-PTB as a template; [**Fig 2L**]) suggested that the canonical and non-canonical modes of binding of K2 to β1-integrin and GIV, respectively, may use two distinct interfaces and remain compatible with the assembly of ternary GIV•K2•β1-integrin complexes in both monomeric and dimeric states of K2 (**Fig S4**). We carried out a series of biochemical assays using recombinant proteins designed to look for intermolecular competition for interacting surfaces and/or the formation of co-complexes *in vitro*. In these assays, two components were kept constant, while the amount of the third component was varied. Increasing His-β1-CT (which contains the NPxY motif recognized by K2) did not displace His-GIV-CT from GST-K2, instead, we unexpectedly observed an enhanced coupling of GIV within a narrow range of concentration of β1-CT, exclusively when His-β1-CT and His-GIV-CT are both present in equimolar concentrations [**Fig 3A**]. Similarly, increasing His-GIV-CT enhanced the coupling of His-K2 to GST-β1-CT only within a narrow range of concentration of His-GIV-CT, exclusively when His-K2 and His-GIV-CT are both present in equimolar concentrations [**Fig 3B**]. Finally, when added in equimolar amounts, His-K2 enhanced the coupling of WT, but not the PGxA mutant His-GIV-CT to GST-β1-CT [**Fig 3C**]. Because these findings were observed consistently, with different protein preparations, the results appear to be most consistent with scenario #3, i.e., GIV and β1-CT may bind K2 non-competitively to assemble ternary complexes when present at optimal stoichiometry [**Fig 3A-C**]. Under the conditions tested, we found that binding of either protein to K2 allosterically augmented the binding of the other only when GIV, β1-CT and K2 are all present in equimolar amounts [**Fig 3A-B**].

**Figure 3.**
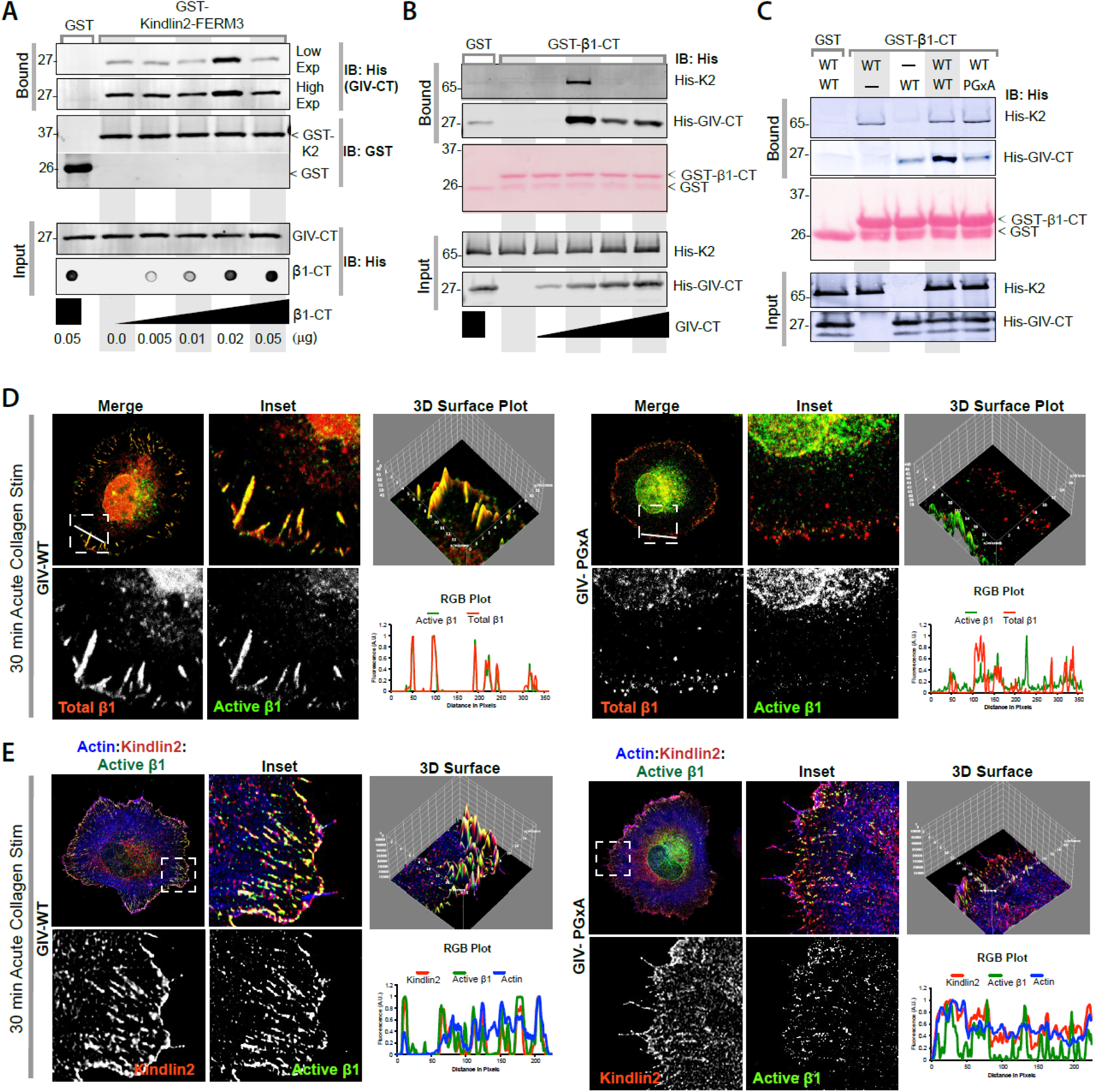
The K2•GIV interaction allosterically augments the K2•β1-integrin interaction and *vice versa*, when present in equimolar proportions, enhances K2 recruitment to focal adhesions and β1-integrin activation. **(A)** GST pulldown assays were performed using fixed concentrations of GST-Kindlin-2-F3-PTB (GST-K2; 0.25 μg) and His-GIV-CT (aa 1660-1870; 1 μg) and increasing concentrations of His-β1 integrin tail, as indicated. Bound GIV was assessed by immunoblotting with anti-His mAb. Equal loading of His-GIV-CT was confirmed by SDS PAGE and the increasing amounts of His-β1 was monitored by dot-blot (input; lower panel). **(B)** GST pulldown assays were performed using equal aliquots of GST-β1 integrin tail (2.5 μg) and full-length His-Kindlin 2 (K2) at equimolar concentrations and increasing amounts of His-GIV-CT (aa 1660-1870). GST-β1-bound proteins were assessed by immunoblotting using anti-His mAb and anti-GIV polyclonal Ab. **(C)** GST pulldown assays were performed using GST-β1 in combination with His-GIV-CT-WT or PGxA (aa 1660-1870) and full-length His-Kindlin 2 (K2). GST-β1-bound proteins were assessed using anti-His mAb and anti-GIV polyclonal Ab. **(D-E)** GIV-depleted MDA-MB-231 cells stably expressing Flag-GIV-WT or PGxA constructs were grown on poly-D-Lysine coated surface, resuspended and plated on collagen coated coverslips for 30 min, fixed and stained for active β1 integrin (green; using a conformational specific antibody, clone 9EG), phalloidin (blue; actin), and either total β1-integrin (D; red) or kindlin-2 (E; red) and analyzed by confocal microscopy. Representative deconvolved images are shown. Insets were analyzed by rendering 3D surface plots and line scans were taken to generate RGB plots using ImageJ.

Next we asked if the allosteric impact of the K2•GIV interaction we observe *in vitro* translates to augmentation of K2•β1-integrin interaction at FAs and β1 activation in cells. To this end, we monitored by immunofluorescence activation of β1-integrins in response to collagen and the recruitment of endogenous K2 to these active receptors in GIV-depleted MDA MB-231 cells stably expressing GIV-WT or PGxA. We found that compared to cells expressing GIV-WT, those expressing GIV-PGxA had reduced integrin clustering and activation, as determined using conformational anti-CD29 9EG7 antibodies [**Fig 3D**]. As expected, with fewer active β1-integrins, the extent of β1•K2 co-localization was also reduced [**Fig 3E**]. Together, these findings indicate that the K2•GIV interaction augments the formation of GIV•K2•β1 complexes and is required for maximal activation of β1-integrins in cells. Such augmentation may be highly regulated by protein stoichiometry and only occur within narrow ranges of protein concentrations. These findings are in keeping with prior observations that much like K2 depletion, K2 overexpression can also cause suppression of β1-integrin activation (Harburger et al., 2009).

### The GIV•kindlin-2 interaction enhances tumor cell adhesion, invasion, integrin signaling

Next we asked how the newly defined GIV•kindlin-2 interaction impact cellular phenotypes. Prior studies have shown that GIV is required for signal amplification downstream of ligand-activated β1-integrins *via* its ability to directly bind and activate Gαi and Class 1 PI3-kinase; the readouts used were cell adhesion and spreading, haptotaxis, activation of the PI3K→Akt, FAK→pY1764GIV and RhoA→myosin light chain (MLC2) signaling axes, the degree of maturation of FAs with sequential recruitment of paxillin and vinculin proteins and activation of FAK (Leyme et al., 2016; Leyme et al., 2015; Lopez-Sanchez et al., 2015). If these downstream events were dependent on GIV’s ability to bind ligand-activated β1-integrins, we hypothesized that uncoupling GIV from β1-integrins will impair them all. Alternatively, if GIV’s ability to trigger G protein and PI3K signaling is independent of its ability to bind and activate β1-integrins, we expected that that uncoupling GIV from β1-integrins will have little or no impact on these readouts. To determine which scenario may be true, we analyzed these readouts in GIV-depleted MDA MB-231 cells (by a shRNA targeting its 3’ UTR [**Fig 4A**]; Lopez-Sanchez et al., 2015) stably expressing GIV-WT and GIV-PGxA that were acutely stimulated by plating on collagen. We found that much like shGIV cells, those expressing the GIV-PGxA mutant was impaired in cell adhesion [**Fig 4A-C**], cell spreading [**Fig 4D**] and haptotaxis along a serum gradient through Matrigel inserts [**Fig 4F-I**]. These phenotypic changes were accompanied also by significant impairment of GIV [**Fig 4J-K; S5A-B**] and Akt phosphorylation [**Fig 4J; S5B**], activation of FAK [**Fig 4L; S5C**] and phosphorylation of MLC2 [**Fig 4M; S5D**]. We used phospho-MLC2 as a surrogate marker of RhoA activity because multiple prior studies (Bhadriraju et al., 2007; Danen et al., 2002; Ren et al., 1999) have shown that RhoA activity drops down to levels below detection during cell adhesion, and hence, monitoring pMLC2 is a more reliable readout of GIV-dependent RhoA activity during integrin signaling (Leyme et al., 2015). Because all these impairments we observed in cells expressing the K2-binding-deficient PGxA mutant of GIV were previously observed in cells without GIV, or those expressing single point mutants of GIV that selectively impair its ability to bind and activate Gαi and PI3K [see **Fig 4M**], we conclude that the GIV•K2 interaction may be an essential upstream event for integrin-coupled downstream activation of Gαi and PI3K. When the interaction is severed (as occurs in cells expressing the PGxA mutant GIV), none of the signaling pathways are effectively triggered, indicating that binding of GIV to integrins within the GIV•K2•β1 complexes is a pre-requisite step.

**Figure 4.**
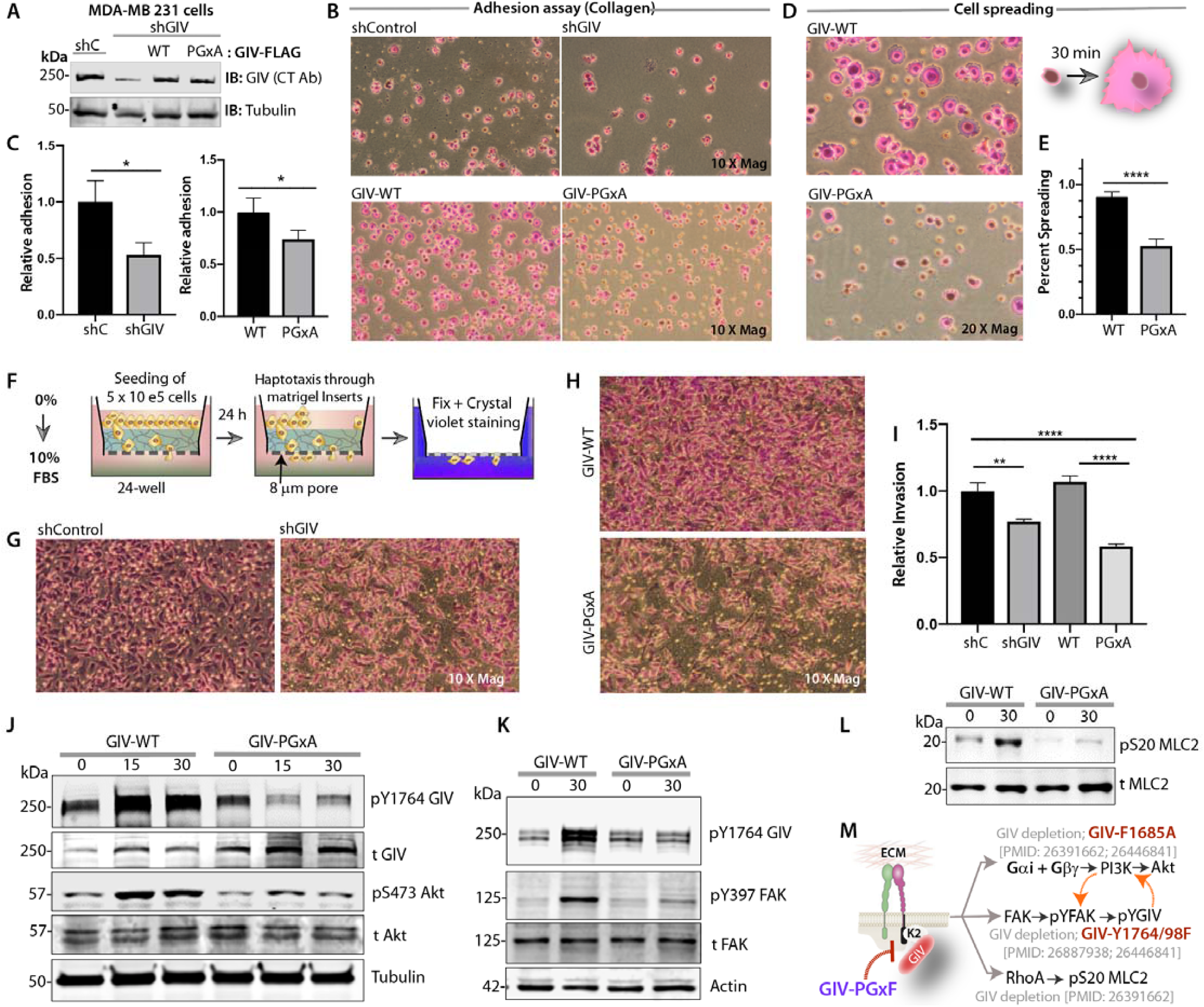
The K2•GIV interaction is required for tumor cell adhesion, spreading and invasion through the ECM. **(A)** Whole cell lysates of control (shC) or GIV-depleted (shGIV) MDA-MB 231 cells stably expressing GIV-WT or the K2-binding deficient PGxA mutant were analyzed for GIV and tubulin by immunoblotting. **(B)** MDA-MB 231 cell lines in A were grown on poly-D-Lysine coated surface, resuspended and plated on 12-well collagen coated plates for 30 min before being fixed in 4% PFA and stained with crystal violet. Cells were visualized and imaged by light microscopy. Representative images are shown. **(C)** Bar graphs display the relative numbers of adherent cells per field, as determined using Image-J. Error bar = S.E.M (n = 4); \**p* = < 0.05 **(D-E)** Adherent MDA-MB 231 cells in B were further analyzed for attachment-induced cell-spreading at higher magnification. Representative images are shown (D) and quantification of % spreading is displayed as bar graph I. Error bar = S.E.M (n = 4); ****p=<0.0001. **(M)** Schematic diagram showing the serum gradient-induced haptotaxis assay conditions used in G-I. **(G-I)** MDA-MB 231 cell lines in A were analyzed for the ability to invade through 26atrigel-coated transwells. The number of cells that successfully invaded within 24 h was imaged (G-H) and quantified using Image-J and displayed as bar graphs (I). Error bar = S.E.M (n = 4); **p=<0.01, ****p=<0.0001. **(J-L)** MDA-MB 231 cells were grown on poly-D-Lysine and stimulated acutely by plating on collagen-coated plates as in B, lysed on plate after indicated periods of time, and equal aliquots of lysates were then analyzed for phospho(p) and total (t) proteins as indicated. See also **Fig S5** for quantification of biologically independent experiments. **(M)** Schematic summarizing the post-receptor pathways previously shown to be impacted by GIV, and the specific approaches (GIV depletion or mutants) that were used to conclude the same. Orange arrows: Positive feed-forward loop of signaling. The mutation F1685A specifically impairs GIV’s ability to bind and activate Gαi (Garcia-Marcos et al., 2009); The mutation Y1764/98F specifically impairs GIV’s ability to bind and activate PI3K (Lin et al., 2011) and enhance the PI3K↔FAK feed-forward loop (Lopez-Sanchez et al., 2015).

Consistent with the observed impairment in integrin signaling, we also found by immunofluorescence/confocal microscopy that the abundance of paxillin and vinculin-positive structures and the activation of FAK in the PGxA-cells were reduced [**Fig 5A**]. To determine if the GIV•K2 interaction persists later in mature FAs, we used two-color super-resolution stimulated emission depletion (STED) microscopy (Hell and Wichmann, 1994; Klar et al., 2000) enabling a lateral resolution of ∼40 nm to assess nanoscale co-localization of endogenous pYGIV and K2 within focal adhesions. Prior studies using STED (Colin-York et al., 2017; Spiess et al., 2018) have revealed the superiority of this approach over conventional microscopy for assessing the organization of FAs into nanometer size clusters of multi-protein assemblies. We observed nanoscale co-localization between pYGIV and K2 within mature FA-structures that were positive for paxillin [**Fig 5B**]; compared to GIV-WT cells, such structures were far fewer and virtually lacking in cells expressing the PGxA mutant. Quantification of these paxillin-positive structures confirmed a significant reduction in the number and size of mature FA-like structures in the PGxA cells.

**Figure 5.**
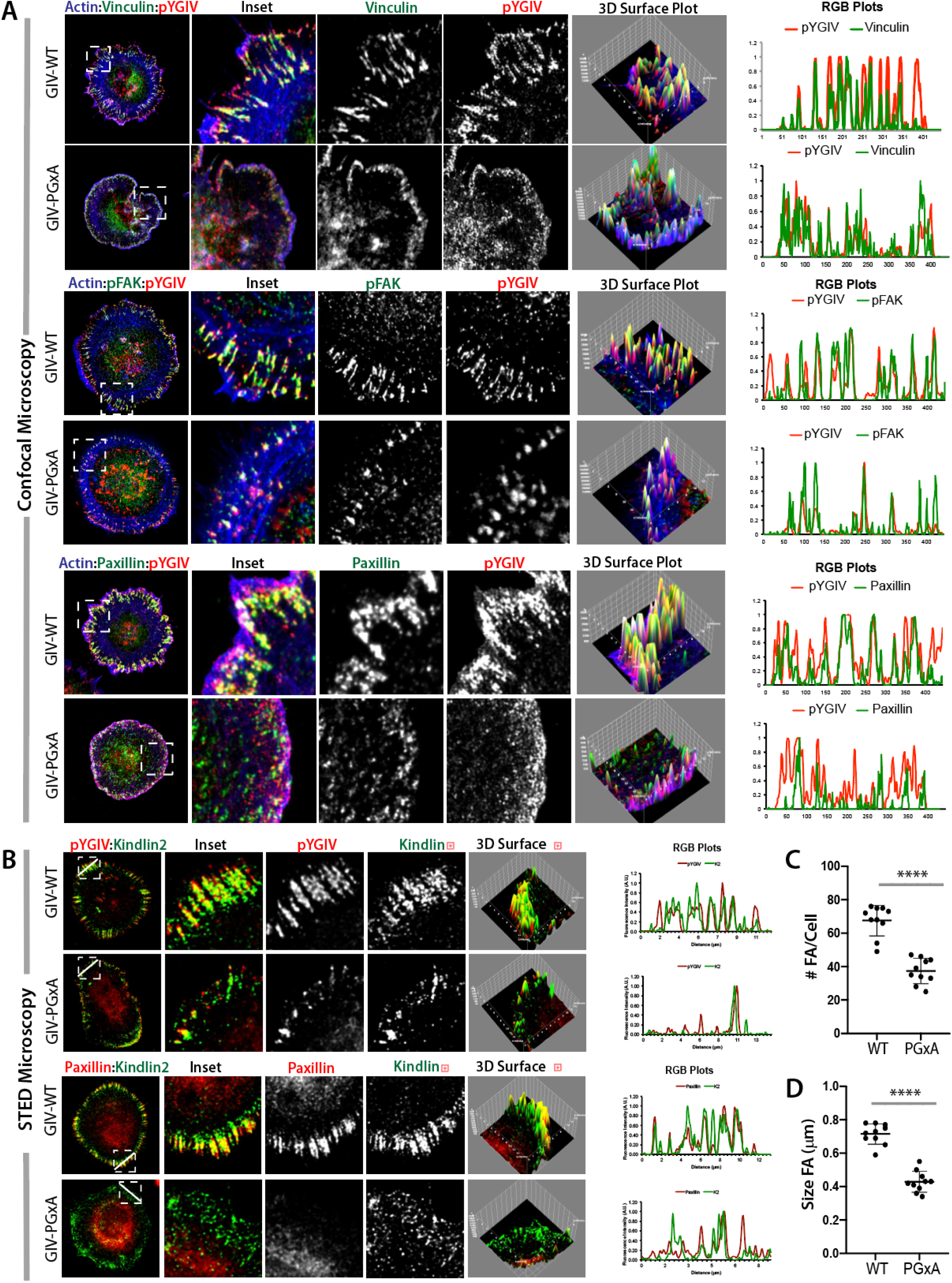
The K2•GIV interaction is necessary for the maturation of focal adhesions (FA), and for triggering the FA-localized FAK→pYGIV signals. **(A)** GIV-depleted MDA-MB-231 cells stably expressing GIV-WT or GIV-PGxA constructs were grown on poly-D-Lysine coated surface, resuspended and plated on collagen coated coverslips for 30 min, fixed and stained for phalloidin [actin; blue], *bona-fide* FA structural or signaling components [i.e., Vinculin (top), pYFAK (Y397; middle) or Paxillin (lower); green], and pYGIV (red) and analyzed by confocal microscopy. Representative deconvolved images are shown. Insets were analyzed by rendering 3D surface plots and line scans were taken to generate RGB plots, both using ImageJ. **(B)** STED super-resolution microscopy was carried out on cells in A to analyze K2 colocalization with pYGIV (top) or paxillin (bottom). Representative deconvolved images are shown. Insets were analyzed by rendering 3D surface plots and RGB plots as in A. **(C-D)** The Paxillin-stained images in A were used to quantify focal adhesion plaque number (C) and size (area; D) with ImageJ.

Taken together, these findings implicate the GIV•K2 interaction in some of the earliest ‘upstream’ events that begin within nascent FAs, i.e., enhanced recruitment of K2 to active β1, formation of GIV•K2•β1-integrin complexes, integrin clustering and activation. Consequently, a myriad of ‘downstream’ events within mature FAs are also derailed, e.g., their number and size, phosphoactivation of MLC, Akt and FAK and the activation of a previously defined feed-forward loop [FAK↔pYGIV↔PI3K↔FAK; (Leyme et al., 2016; Lopez-Sanchez et al., 2015)]. It is certainly possible that some of the post-receptor activation pathway/downstream signaling in PGxA mutant cells may reflect some of the impact of GIV’s ability to bind the PTB domain of Tensin. However, the observed impact of GIV-PGxA on integrin activation observed earlier (**Fig 3D-E**) is unlikely to be due to GIV•Tensin interaction because tensin is recruited much later (Torgler et al., 2004) than the time points studied here, in mature focal adhesions (unlike GIV, which colocalizes with β1-integrin in nascent adhesions at the cell periphery; (Lopez-Sanchez et al., 2015)), has been shown to be a part of downstream signaling pathway (Bockholt and Burridge, 1993) and is considered as largely dispensable for the early steps of integrin activation (Calderwood et al., 2013; Theodosiou et al., 2016).

### The GIV•kindlin-2•β1-integrin synergy may have a poor prognosis in breast cancer

Next we asked if the observed allosteric synergy between GIV, K2 and β1-integrin we observe here and the impact of such synergy on sinister properties of tumor cells can be meaningful when assessing tumor behavior and/or prognosticating clinical outcome. First we asked if the levels of expression of each of these three entities alone, or in combination could impact one of the most important readouts of cancer aggressiveness, i.e., metastasis-free patient survival. To discern this, we chose to study a pooled cohort of patients (Bos et al., 2009; Minn et al., 2005; Wang et al., 2005) with breast cancers who did not receive adjuvant chemotherapy, and hence, in them metastatic progression reflects natural disease progression and not resistance/selection under treatment. Samples were divided into “low” and “high” subgroups with regard to GIV (CCDC88A; **6A**), β1-integrin (ITGB1; **6B**) and K2 (FERMT2; **6C**) gene expression levels using the StepMiner algorithm (Sahoo et al., 2007), implemented within the hierarchical exploration of gene-expression microarrays online (HEGEMON) software (Dalerba et al., 2011; Volkmer et al., 2012). Kaplan-Meier analyses of the disease free-survival showed that high levels of expression of each alone was enough to carry a grave prognosis, i.e., a shorter metastasis-free survival [**Fig 6A-C**]. To determine if the levels of expression of these genes, and a few other important genes implicated in some of the earliest steps of integrin activation interact synergistically as independent variables to impact survival of patients, we carried out a Cox proportional-hazards model (Cox, 1972) which is a regression model that is commonly used as a statistical method for investigating the effect of several variables upon the time it takes for a specified event to happen, in this case, metastasis (Cox, 1972). We found that in this model, CCDC88A (GIV) significantly interacts with FERMT2 (K2), ITGB1 (β1-integrin), TNS1, 4 (tensin), but not TLN1 (talin); FERMT2 (K2) only interacts with GIV. ITGB1 interacted with GIV and tensin; talin interacted primarily with ITGB1. Thus, of all the gene pairs tested, high GIV and K2 expression had a synergistic impact on shortening metastasis-free survival. These findings not only provide evidence for ‘interaction’ between GIV and K2 at the levels of gene expression and expose its impact on survival outcome, but also serves to further cement the role of the GIV•K2 interaction in tumor aggressiveness and progression to metastasis. Findings are also consistent with prior evidence that elevated levels of K2 carries a poor prognosis in a variety of cancers (Cao et al., 2015; Ge et al., 2015; Ma et al., 2015b; Mahawithitwong et al., 2013; Shen et al., 2012; Sin et al., 2011; Talaat et al., 2011; Yan et al., 2016; Zhan et al., 2015),

**Figure 6.**
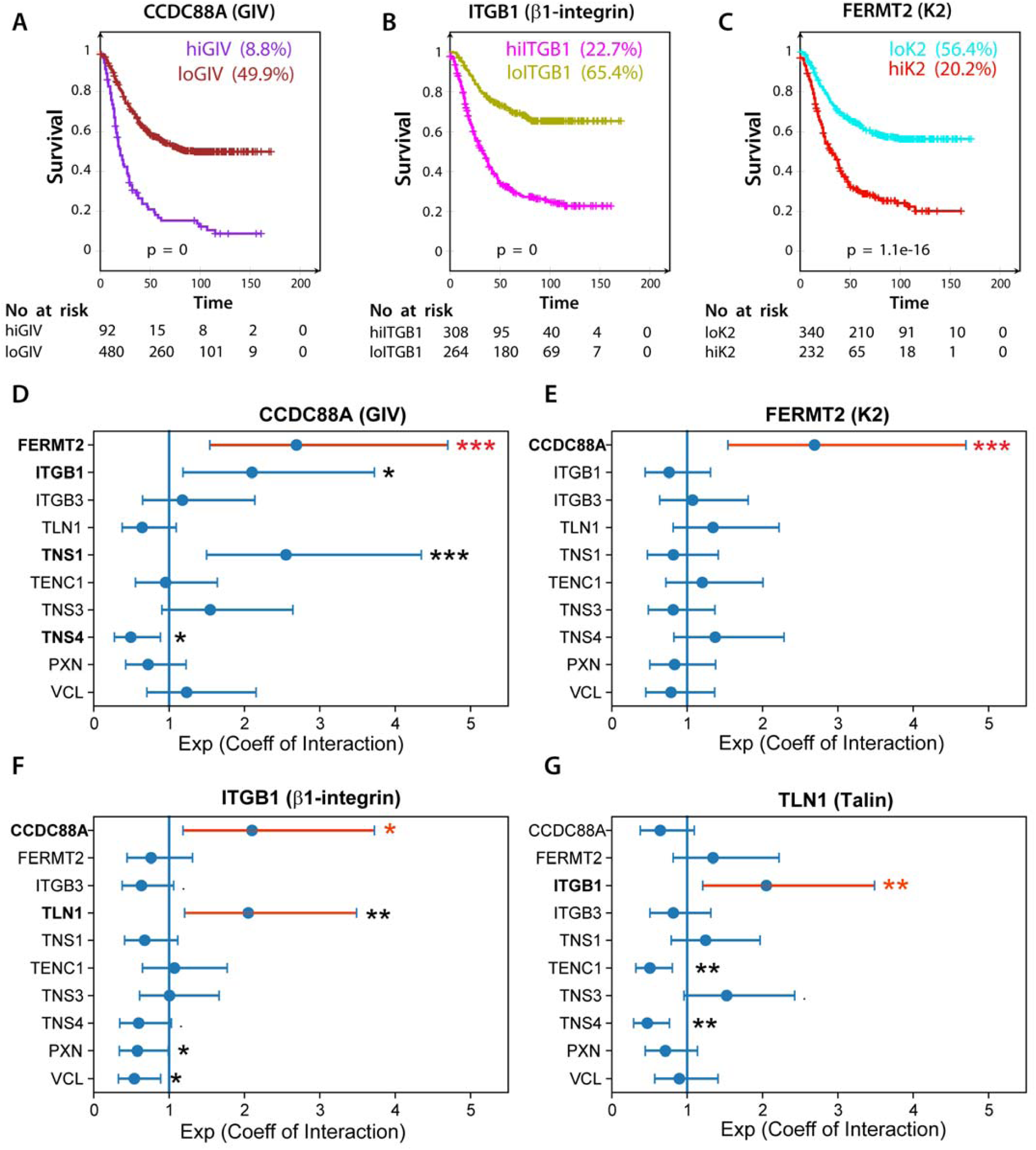
Levels of expression of GIV, Kindlin-2 and β1-integrin prognosticate progression to metastasis. Primary breast cancers from 572 patients (3 independent cohorts, pooled (Bos et al., 2009; Minn et al., 2005; Wang et al., 2005)) who did not receive adjuvant therapy were segregated into groups of high vs low expression (see *Methods*) of target genes and analyzed for metastasis-free survival. **(A-C)** Kaplan–Meier curves for metastasis-free survival over time among patients whose primary tumors had high vs low levels of GIV (CCDC88A; A), β1-integrin (ITGB1; B) and K2 (FERMT2; C). **(D-G)** Statistical interaction (synergy between variables) is measured in the Cox proportional hazards regression model for CCDC88A (GIV; **D**), FERMT2 (K2; **E**), ITGB1 (β1-integrin; **F**) and TLN1 (talin; **G**) with other major genes involved in some of the earliest steps of integrin activation and the formation and maturation of the FAs. Coefficient of the interaction term in Cox regression models is plotted with 95% confidence interval that demonstrates the significance of the statistical interaction. *p* values: ‘***’ 0.001 ‘**’ 0.01 ‘*’ 0.05.

## CONCLUSIONS

The major discovery we report herein is the mechanism and consequences of a direct interaction between GIV and K2. Using selective single point mutants of GIV that are incapable of binding K2, we chart the two major consequences of this GIV•K2 interaction-- First, the GIV•K2 interaction impacts GIV biology because it appears to be a pre-requisite for the previously defined GIV-dependent signaling programs downstream of ligand-activated integrins, e.g., G protein (Gβγ→PI3K), PI3K→Akt, PI3K→FAK→pYGIV and RhoA→MLC. Second, this interaction impacts integrin biology because it augments the affinity of K2 for the cytoplasmic tail of β1-integrins *in vitro*, the recruitment of K2 to β1-integrins in cells and its subsequent clustering and activation within nascent and mature FAs.

As for the impact of the GIV•K2 interaction on GIV biology, findings showcased here, together with our understanding of how GIV modulates integrin/FAK signaling via G protein intermediates (Leyme et al., 2016; Leyme et al., 2015; Lopez-Sanchez et al., 2015), provide a more complete mechanistic insight into the roles of GIV in both early and late steps of integrin signaling [**Fig 7A**]. Because integrin signaling aids cancer growth, metastasis, and drug resistance, the signaling interfaces assembled by GIV (GIV•K2 and GIV•Gαi) provide new strategies for targeting the integrin pathway. Furthermore, GIV’s FERM3/PTB-binding PGxF motif provides the 2^nd^ mechanism [GIV’s SH2-like module was the 1^st^(Lin et al., 2014)] by which GIV can couple Gα-proteins to non-GPCRs like integrins [**Fig 7B**] which are typically believed to initiate tyrosine-based signals. Because GIV binds the PTB-domain of tensin, it is possible that the newly defined SLIM in GIV that binds K2-FERM3-PTB and tensin-PTB may also recognize other PTB-module containing adaptors. If so, it is possible that such versatility of the motif could enable coupling of GIV/G proteins, PI3K activation and RhoA-dependent actin remodeling to diverse classes of non-GPCRs besides integrins that also use PTB proteins as adaptors, i.e., cytokine, LDL-receptor, leukocyte receptors, RTKs, and others (Smith et al., 2006), leading to signal convergence.

**Figure 7.**
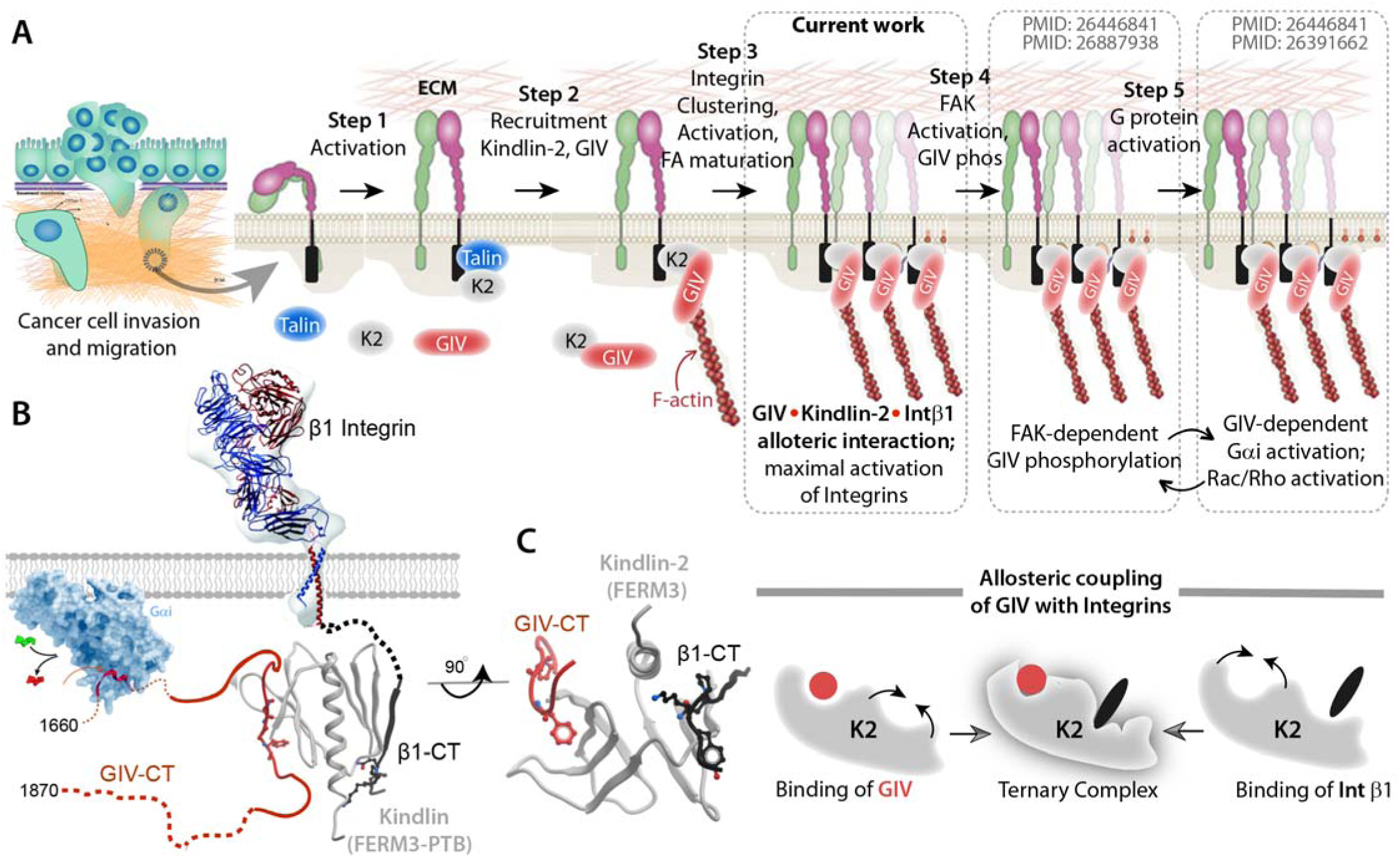
Schematic summarizing how GIV impacts early and late events in β1-integrin signaling. **(A)** Schematic illustrating the role of GIV during various steps of β1-integrin activation and signaling. Integrin activation is mediated by binding of its cytoplasmic tail to Talin (Step 1) and GIV•K2 complexes (Step 2). The latter is necessary for integrin clustering, activation and maturation of focal adhesions (Step 3), activation of FAK and tyrosine phosphorylation of GIV (Step 4). Previously described forward feedback loops (Step 5) orchestrated by G protein signaling further enhances integrin signaling. **(B)** Structural basis for how ligand-activated β1-integrins may bind and modulate trimeric G protein, Gαi via the assembly of K2•GIV complexes. **(C)** Binding of either GIV or β1-integrin on non-overlapping interfaces of K2 allosterically enhances the formation of GIV•K2•β1-integrin complexes.

As for the impact of the GIV•K2 interaction on integrin biology, its ability to allosterically augment the affinity of K2 for β1-integrins and enhance the activation of β1-integrins resembles how talin-dependent activation of β1-integrins is triggered by adjusting the affinity of talin for integrins. For example, binding of talin’s FERM domain to charged acidic phospholipids—phosphatidylinositol 4,5-bisphosphate(PtdIns(4,5)P2) greatly increased its affinity for integrins, so that once talin is recruited to the plasma membrane, this phospholipid could augment the formation of talin–integrin complexes (Moore et al., 2012; Ye et al., 2016). In fact, membrane recruitment of talin is sufficient to increase its affinity for integrins (Han et al., 2006; Lagarrigue et al., 2016; Lee et al., 2009). In the case of K2, researchers generally agree that K2’s adaptor functions may be critical for maximal integrin activation and clustering [reviewed in (Calderwood et al., 2013)], but how the K2•β1-integrin interaction augments integrin activation remained unknown (Bledzka et al., 2012; Kahner et al., 2012). We have not only pinpointed GIV as one binding partner that adjusts the affinity of K2 for β1-integrins, but also provided a molecular basis for how such adjustment of affinity is brought about through an intramolecular allostery within K2 [**Fig 7C**]. Our findings that GIV and β1-integrin may augment each other’s ability to bind K2-FERM3-PTB suggest that the interplay between integrin, K2 and GIV may serve as one of the long-sought missing early steps in integrin activation.

## Acknowledgements

This paper was supported by NIH CA238042, AI141630, CA100768 and CA160911 (to P.G). C.R was supported by an NCI/NIH-funded Cancer Therapeutics Training (CT^2^) Training Program (T32 CA121938) and a NIDDK/NIH-funded training grant (T32 DK007202). N.A.K was supported by a NIH predoctoral fellowship (F31 CA206426), and T32 training grants T32CA067754 and T32DK007202. S.R. was supported by NIH R01-- AI118985 and GM117424 (to Handel and Kufareva) and in part by an intramural UC San Diego Chancellor’s Center Launch Funds. We thank Ying Dunkel and Nina Sun, for technical support in this work. We also thank Dr. Irina Kufareva at Skaggs School of Pharmacy and Pharmaceutical Sciences for insightful discussions during the work.

## Author Contributions

C.R, N.A.K, N.R and P.G designed, executed and analyzed most of the experiments in this work. S.R cloned and purified His-Kindlin and GST-kindlin constructs and both I.L-S and S.R performed pulldown studies with WT and mutant Kindlin constructs. C.R, N.A.K and P.G conceived the project. C.R and S.R wrote materials and methods and edited the manuscript. P.G wrote the manuscript.

