## Supplementary Online Materials for "GIV•Kindlin interaction is required for Kindlin-Mediated Integrin Recognition and Activation"

### INVENTORY OF SUPPLEMENTARY MATERIALS

- **STAR METHODS**
- **SUPPLEMENTARY FIGURES AND LEGENDS**
- **BIBLIOGRAPHY**

**STAR *Methods**

- **Key Resource Table**
- **Contact for Reagent and Resource Sharing**
- **Experimental Model and Subject Details**
    - Cell Lines (Cos7, MDA-MB-231)
- **Method Details**
    - Cell culture
    - Whole Cell Immunofluorescence
    - Western Blotting
    - In Vitro Pulldown and Co-immunoprecipitation (CoIP)
    - Recombinant Protein Purification
    - Adhesion Assay
    - Transwell Invasion Assay
- **Quantification and Statistical Analysis**
    - Statistical Analysis
    - Replications
- **Data and Software Availability**

**Key Resource Table:**

| REAGENT or RESOURCE | SOURCE | IDENTIFIER |
|---|---|---|
| | **ANTIBODIES** | |
| **Rabbit monoclonal anti-pY1765 GIV** | Roche Spring Biosciences | 06974937001 Clone SP158 |
| **Rabbit polyclonal anti-GIV CT** | Santa Cruz Biotechnology | N/A |
| **Rabbit polyclonal anti-G$\alpha$i3 (C-10)** | Santa Cruz Biotechnology | N/A |
| **Rabbit polyclonal anti-$\beta$-tubulin** | Santa Cruz Biotechnology | sc-9104 |
| **Mouse monoclonal anti-GAPDH** | Santa Cruz Biotechnology | sc-365062 |
| **Mouse monoclonal anti-HIS** | GenScript | A00186-100 |
| **Mouse monoclonal anti-GST** | GenScript | A00865 |
| **Mouse monoclonal anti-$\alpha$-tubulin** | Santa Cruz Biotechnology | sc-5286 |
| **Mouse monoclonal anti-pFAK** | BD Biosciences | 611722 |

| | | |
|---|---|---|
| **Mouse monoclonal anti-paxillin** | BD Biosciences | 610051 |
| **Mouse monoclonal anti-vinculin** | Sigma-Aldrich | V9131 |
| **Rabbit polyclonal anti-paxillin** | Santa Cruz Biotechnology | SC-5574 |
| **Mouse monoclonal anti-kindlin2** | EMD Millipore | MAB2617 |
| **Rat anti-active β1 integrin** | BD Pharmingen | 9EG7 |
| **Mouse monoclonal anti-β1 integrin** | Abcam | Ab30394 |
| **Goat anti-Rabbit IgG (680)** | LI-COR Biosciences | 926-68071 |
| **Goat anti-Rabbit IgG, Alexa Fluor 594 conjugated** | ThermoFisher Scientific | A11072 |
| **Goat anti-Rabbit IgG, Alexa Fluor 647 conjugated** | ThermoFisher Scientific | A27040 |
| **Goat anti-Mouse IgG (800)** | LI-COR Biosciences | 926-32210 |
| **Goat anti-Mouse IgG, Alexa Fluor 488 conjugated** | Thermo Fisher Scientific | A11017 |
| **Biological Samples and Cell Lines** | | |
| **Cos7** | ATCC | CRL-1651 |
| **MDA-MB-231** | ATCC | HTB-26 |
| **CHEMICALS, RECOMBINANT PROTEINS, AND PLASMIDS** | | |
| **DAPI (4',6-Diamidino-2-Phenylindole, Dilactate)** | Thermo Fisher Scientific | D3571 |
| **Collagen I** | BD Biosciences | 354249 |
| **Poly-D-lysine** | Millipore Sigma | A-003-E |
| **Mirus TranIT LT1** | Mirus Bio LLC | MIR2300 |
| **G418** | Cellgro | A-1720 |
| **Puromycin** | Life Technologies | A1113803 |
| **Biocoat Matrigel Invasion Inserts** | Corning | 354480 |
| **Paraformaldehyde 16%** | Electron Microscopy Biosciences | 15710 |
| **Phalloidin 594** | Thermo Fisher Scientific | A12381 |
| **Prolong Gold** | Thermo Fisher Scientific | P10144 |
| **Prolong Glass** | Thermo Fisher Scientific | P36980 |
| **pGEX-4T-GIV-CT-WT (a.a. 1623-1870)** | (Garcia-Marcos et al., 2009) | N/A |
| **pET-28b-GIV-CT-WT (a.a. 1660-1870)** | (Garcia-Marcos et al., 2009) | N/A |
| **pET-28b-GIV-CT-PGxA(a.a. 1660-1870)** | This paper | N/A |
| **CMV14-p3X FLAG-GIV WT (full length)** | This paper | N/A |
| **pGEX4T-Kindlin C term ; [GST-mFERM2 (571-680)]** | (Montanez et al., 2008) | N/A |

| | | |
|---|---|---|
| **pGEX-4T1-Kindlin2 C-term QW/AA [GST-mFERM2 (571-680), Q614A/W615A]** | (Montanez et al., 2008) | N/A |
| **GST-mFERM2(564-680)** | This paper | N/A |
| **GST-mFERM2(564-680) QW/AA [Q614A/W615A]** | This paper | N/A |
| **6xHis-SUMO-Kindlin2Δ** | (Li et al., 2017) | N/A |
| **CMV14-p3X FLAG-GIV PGxA (full length)** | This paper | N/A |
| **pGEX-4T1-Tensin3-SH2-PTB** | (Qian et al., 2009) | N/A |
| **pGEX-4T1-Tensin3 -SH2** | (Qian et al., 2009) | N/A |
| **pGEX4T-Integrin beta 3** | (Montanez et al., 2008) | N/A |
| **pGEX4T-Integrin beta 1** | (Montanez et al., 2008) | N/A |
| **pGEX-6P-GST- Talin-F3 domain** | (Ye et al., 2010) | N/A |
| **Kindlin2 siRNA** | Santa Cruz Biotechnology; (Yang et al., 2016) | sc-106786 |
| **SOFTWARE** | | |
| **ImageJ** | National Institute of Health | https://imagej.net/Welcome |
| **Prism** | GraphPad | https://www.graphpad.com/scientific-software/prism/ |
| **LAS-X** | Leica | www.leica-microsystems.com/products/microscope-software/p/leica-las-x-ls |
| **Molsoft ICM v3.8-6** | Molsoft LLC | http://www.molsoft.com/index.html |

### Methods

#### Plasmid constructs and protein expression

Cloning of GIV-CT (aa 1660–1870) into pET28b (His-GIV CT) was described previously (Garcia-Marcos et al., 2012) GST-K2F3, GST-β1-CT and GST-β3-CT plasmids were generously obtained from Reinhardt Fassler (Max Planck Institute, Germany). GST-Tensin3 plasmid was generously obtained from Douglas Lowy, the GST-Talin plasmids and His-β1-CT plasmids were obtained from Mark H. Ginsberg (UC San Diego), and the 6xHis-SUMO-Kindlin2Δ construct (Li et al., 2017) was generously provided by Prof. Cong Yu (.Southern University of Science and Technology, Shenzhen, China). For mammalian expression, RNA interference–resistant (shRNA rest) GIV was cloned into p3XFLAG-CMVTM-14 plasmid (GIV-FLAG) as described previously (Garcia-Marcos et al., 2009) GIV-FLAG and His-GIV-CT mutants (F1742A) were generated by site-directed mutagenesis using a QuikChange kit (Stratagene, CA, USA) and specific primers (sequence available upon request) as per the manufacturer's protocols (Garcia-Marcos et al., 2009; Lin et al., 2011) Primer sequences are available upon request. shRNA 3–untranslated region for GIV (GIV shRNA: CCGGGCTTTCATT-ACCAGCTCTGAACTCGAGTTCAGAGCTGGTAATGAAAGCTTTTTTG) was cloned into pLKO.1 (TRCN0000130452) or control vector TRC1.5-pLKO.1-puro. His-GIV-CT fusion construct was expressed in E. coli strain BL21 (DE3) and purified as described previously (Garcia-Marcos et al., 2009; Ghosh et al., 2008). Briefly, bacterial cultures were induced overnight at 25°C with 1 mM isopropyl β-d-1-thiogalactopyranoside (IPTG). Pelleted bacteria from 1 l of culture were resuspended in 10 ml of His lysis buffer (50 mM NaH2PO4, pH 7.4, 300 mM NaCl, 10 mM imidazole, 1% [vol:vol] Triton X-100, 2X protease inhibitor cocktail [Complete EDTA-free, Roche Diagnostics, CA, USA]). After sonication (3 × 30 s), lysates were centrifuged at 12,000 × g at 4°C for 20 min. Solubilized proteins were affinity purified on HisPur Cobalt Resin (Pierce, IL, USA). Proteins were eluted, dialyzed overnight against PBS, and stored at −80°C.

GST-mFERM2(564-680) (encoding for the F3 domain of mouse Kindlin2, Uniprot Q8CIB5, C-terminal to GST and the SDLVPRGSPEL linker, as part of a pGX4T-1 expression vector) was generated from the GST-mFERM2(571-680) generously shared by Reinhard Faessler (Max Planck Institute, Germany) (Montanez et al., 2008) Similarly, GST-mFERM2(564-680) Q614A/W615A was made from GST-mFERM2(571-680) Q614A/W615A. The constructs were generated using standard site directed mutagenesis protocol. The PCR products were amplified using the forward primer: TCTCTGCCTGAGTTCGGCATCACACACTTCATTGCGAGG and reverse primer: GATGCCGAACTCAGGCAGAGACAATTCCGGGGATCCACG, using Pfu Turbo (Agilent, CA, USA), digested by *Dpn I* (NEB, UK), transformed into XL10 (Gold) competent cells, and grown on agar plates with 50 mg/mL Carbenicillin. All the aforementioned constructs were checked by sequencing with PGEX universal primer (Genewiz, USA).

#### *Recombinant Protein Purification*

Both GST and His-tagged proteins were expressed in E. coli stain BL21 (DE3) and purified as previously described (ref). Briefly, cultures were induced using 1mM IPTG overnight at 25°C. Cells were then pelleted and resuspended in either GST lysis buffer (25 mM Tris-HCl, pH 7.5, 20 mM NaCl, 1 mM EDTA, 20% (vol/vol) glycerol, 1% (vol/vol) Triton X-100, 2×protease inhibitor cocktail) or His lysis buffer (50 mM $NaH_2PO_4$ (pH 7.4),

300 mM NaCl, 10 mM imidazole, 1% (vol/vol) Triton X-100, 2X protease inhibitor cocktail). Cells were lysed by sonication, and lysates were cleared by centrifugation at 12,000 X g at 4°C for 30 mins. Supernatant was then affinity purified using glutathione-Sepharose 4B beads (GE Healthcare) or HisPur Cobalt Resin (Thermo Fisher Scientific), followed by elution, overnight dialysis in PBS, and then storage at -80°C.

For expressing and purifying His-Kindlin-2, 6xHis-SUMO-Kindlin2Δ plasmid was transformed into Rosetta cells for protein expression. For large scale purification, 2 L of secondary culture was induced at 25°C overnight using 0.2 mM IPTG. Cell pellet was resuspended in Resuspension Buffer (RB) consisting of 50mM phosphate buffer pH 7.4, 300mM NaCl, 5 mM bME, 5 mM imidazole, DNAse, and 1 tablet of protease inhibitor cocktail (Roche, added fresh before use). The suspension was sonicated (Branson Digital sonicator) with 18% pulse with 30 second time on and off for five minutes each until the protein suspension was clear, after which Triton X-100 was added to 0.1%. The suspension was centrifuged at 13,000 rcf for 45min. The supernatant was collected and incubated with Talon resin (Takara, Japan) for 2h at 4°C. The resin was then washed with RB + 50 mM imidazole and eluted with RB + 200 mM imidazole. The eluted protein was loaded into a size exclusion column (Superdex 200 Increase 10/300, GE Healthcare, USA) with SEC buffer as HEPES pH 7.5, 100 mM NaCl, 1mM $MgCl_2$, 1 mM DTT. The fractions were collected, pooled, spin-concentrated using Amicon® Ultra (Sigma, MO, USA) with a 30 kDa MW cutoff, flash-frozen in small aliquots, and stored at -80 for further use.

***In Vitro Pulldown and Co-immunoprecipitation (Co-IP)***

Purified GST-tagged proteins from E. coli were immobilized onto glutathione-Sepharose beads and incubated with binding buffer (50 mM Tris-HCl pH 7.4, 100 mM NaCl, 0.4% (v:v) Nonidet P-40, 10 mM $MgCl_2$, 5 mM EDTA, 2 mM DTT, 1X Complete protease inhibitor) for 60min at room temperature. For GST-pulldown assays with recombinant proteins, the proteins were diluted in binding buffer and incubated with immobilized GST-proteins for 90min at room temperature. For binding with cell lysates, cells were lysed in cell lysis buffer (20 mM HEPES pH 7.2, 5 mM Mg-acetate, 125 mM K-acetate, 0.4% Triton X-100, 1 mM DTT, 500 µM sodium orthovanadate, phosphatase inhibitor cocktail (Sigma-Aldrich, MO, USA) and protease inhibitor cocktail (Roche Life Science)) using a 28G syringe, followed by centrifugation at 10,000Xg for 10min. Cleared supernatant was then used in binding reaction with immobilized GST-proteins for 4 hours at 4°C. After binding, bound complexes were washed four times with 1 ml phosphate wash buffer (4.3 mM $Na_2HPO_4$, 1.4 mM $KH_2PO_4$, pH 7.4, 137 mM NaCl, 2.7 mM KCl, 0.1% (v:v) Tween 20, 10 mM $MgCl_2$, 5 mM EDTA, 2 mM DTT, 0.5 mM sodium orthovanadate). Bound proteins were then eluted through boiling at 100°C in Laemmli buffer (BIORAD, CA, USA). For experiments using Kindlin-2 constructs (both GST and His), the bound proteins were eluted at 37°C for 10 min.

***Cell culture, transfection, lysis, and quantitative immunoblotting***

Cos7, HeLa and MDA-MB-231 cells were obtained from American Type Culture Collection (ATCC). Transfection, lysis, and immunoblotting were carried out exactly as described before (Aznar et al., 2016; Lopez-Sanchez et al., 2015) Cells were transfected using Mirus LT1 following the manufacturers' protocols. For assays involving serum starvation, serum concentration was reduced to 0% FBS overnight.

Whole-cell lysates were prepared after washing cells with cold PBS before resuspending and boiling them in sample buffer. Lysates used as a source of proteins in pull-down assays were prepared by resuspending cells in lysis buffer (20 mM HEPES, pH 7.2, 5 mM Mg acetate, 125 mM K acetate, 0.4% Triton X-100, and 1 mM dithiothreitol supplemented with sodium orthovanadate [500 μM], phosphatase [Sigma-Aldrich, MO, USA], and protease [Roche, USA] inhibitor cocktails), after which they were passed through a 28-gauge needle at 4°C and cleared (10,000 × g for 10 min) before use in subsequent experiments.

For immunoblotting, protein samples were separated by SDS-PAGE and transferred to polyvinylidene fluoride membranes (Millipore Sigma, MO, USA). Membranes were blocked with PBS supplemented with 5% nonfat milk (or with 5% BSA when probing for phosphorylated proteins) before incubation with primary antibodies. In some instances, the samples were separated on a 12% SDS PAGE and transferred to a nitrocellulose membrane (Bio-Rad, CA, USA) using TransBlot-Turbo (Bio-Rad, CA, USA). The membrane was stained with Ponceau S to visualize baits, then washed and blocked with PBS with 0.1% Tween (PBS-T) and 0.5% BSA overnight at 4°C. Infrared imaging with two-color detection and quantification were performed using a Li-Cor Odyssey imaging system. Dilution of primary antibodies used were as follows: anti–GIV-CT, 1:500; anti-Gαi3, 1:333; anti-β tubulin, 1:1000; anti-β1 integrin, 1:250; and anti–His, 1:500. All blots were visualized using LI-COR Odyssey imager, and band analysis was performed with Image Studio™ Lite 5.2 (LI-COR Biosciences, NE, USA). Figures were assembled for presentation using Photoshop (Adobe, San Jose, CA, USA) and Illustrator (Adobe, San Jose, CA, USA) software.

#### ***Generation of stable cell lines***

shRNA control and shRNA GIV MDA-MB-231, stable cell lines using Mission RNAi technology (Sigma-Aldrich, MO, USA) were generated by lentiviral transduction followed by selection with puromycin (2.5 μg/ml) a described previously (Midde et al., 2018). Depletion of endogenous GIV was confirmed by immunoblotting with GIV-CT rabbit antibody. Lentiviral packaging was performed in HEK293T cells by co-transfecting the shRNA constructs with psPAX2 and pMD2G plasmids (4:3:1 ratio, respectively), using Mirus LT1. The medium was changed after 24 h, and virus-containing medium was collected after 36–48 h and centrifuged and filtered through a 0.45-μm filter. psPAX2 and pMD2G plasmids were a generous gift from Christopher K. Glass (University of California, San Diego, La Jolla, CA). shRNA GIV MDA-MB-231 stable cell lines expressing p3xFLAG-CMV-14-GIV (GIV-3xFLAG) WT and PGxA constructs were selected as previously described (Aznar et al., 2016) with the neomycin analogue G418 at 800 μg/ml. Expression of various GIV constructs were confirmed to be similar to levels of endogenous GIV in shRNA control cells by immuno-blotting with GIV-CT antibodies.

#### ***Whole-cell confocal immunofluorescence***

Cells were fixed at room temperature with 3% PFA in PBS for 15 min, treated with 0.1 M glycine for 10 min, and subsequently permeabilized for 1 h (0.2% Triton X-100 in PBS) and blocked in PBS containing 1% bovine serum albumin (BSA) and 0.1% Triton X-100 as described previously (Lopez-Sanchez et al., 2014). Primary and secondary antibodies where incubated for 1h at room temperature in blocking buffer. ProLong Gold or Prolong Glass (Life Technologies, USA) was used as mounting medium. Dilutions of antibodies used were as follows: phosphotyrosine (pY)-1764-GIV, 1:300; vinculin, 1:400; paxillin, 1:200; integrin-β1, 1:400; phalloidin, 1:1000; phospho-FAK, 1:100; kindlin-2, 1:150; conformational specific antibody integrin-β1, 1:400; DAPI, 1:2000; and secondary goat anti-rabbit (488), goat anti-mouse (594), and goat-anti-mouse or rabbit (647) Alexa-conjugated antibodies, 1:500. Images were acquired at room temperature with a Leica TCS SPE-II with DMI4000 microscope equipped with a Leica Hamamatsu 9100-02 camera and the LAS AF SPE software (Leica, Germany) using a 63× oil-immersion objective using 488-, 561-, 635-, and 405-nm laser lines for excitation. The settings were optimized, and the final images scanned with line averaging of three scans. Images were further processed using Lightning deconvolution in the Leica LAS-X (Leica Microsystems, Germany) software package.

Quantification of focal adhesions was carried out using the particle analyzer application on ImageJ (NIH, MD, USA) exactly as outlined previously (Horzum et al., 2015 blue right-pointing triangle). All images were processed using ImageJ software and assembled for presentation using Photoshop and Illustrator software. Images shown are representative of 90–95% cells that were evaluated across three independent experiments.

***Molecular modeling***

A NMR-resolved structure of DLC1(peptide) bound to the PTB domain of tensin [(Chen et al., 2012) PDB 2loz] and a structure of β1-integrin-bound dimerized 6xHis-SUMO-Kindlin2Δ [(Li et al., 2017) PDB 5xq0] were used as docking templates to model β1-integrin-bound Kindlin-2. Modeling was carried out using Molsoft (see STAR materials Table).

***Super Resolution* Stimulated emission depletion (STED*) microscopy***

This was performed using a Leica TCS SP8 Confocal microscope equipped with a white light laser tunable to excitations between 470 and 670nm. Cells were plated on collagen-coated coverslips and allowed to adhere for 30 min. They were then fixed with 4% PFA and stained with antibodies against kindlin2 (1:100), pYGIV (1:100) or Paxillin (1:100) followed by secondary antibody incubation (AlexaFluor 488 and AlexaFluor 647 1:500). Coverslips were then mounted on Prolong Glass and allowed to cure for 5 days before imaging. Images were acquired using the HyD detectors with the tunable white light laser set to the corresponding Alexa 488 and Alexa 647 nm excitation and emission wavelengths. Because of the combination of fluorophores used depletion lasers of 592 nm were used for the 488 channel and depletion laser of 775 nm used for the 647 nm channel. A 100x oil objective was used to acquire all images. Once images were acquired, they were deconvolved using LAS-X software and then processed using ImageJ. All images were further processed in ImageJ using the 3D surface plot plugin and line scans were done to generate RGB plots.

***Transwell invasion assay***

Cell invasion was assessed using Biocoat Matrigel (Corning, NY, USA) inserts with 8-μm pores in 24-well plates. Cells were detached using trypsin/EDTA and resuspended in DMEM supplemented with 0.4% FBS. A total of 5 × 105 cells was loaded in the upper well in a volume of 300 μl, and the lower well was filled with 750 μl of DMEM with 10% FBS. The plates were incubated at 37°C for 24h before removing the remaining cell suspension. The invasion insert was placed in a clean well containing 4% PFA for 1 h at room temperature, stained with crystal violet for 1 h, and washed three times in PBS. Cells on the upper side of the filters were removed with cotton-tipped swabs, and the number of migrated cells on the bottom side of the filter was counted in five randomly chosen fields at 200× magnification and averaged. All experiments were performed in triplicate, and each experiment was repeated at least three times.

***Adhesion Assay***

12-well plates were coated with collagen, rinsed with PBS, and then blocked with 0.5% BSA for 1h at 37°C. Cells were harvested with trypsin/EDTA, seeded in the wells at 2 × 104 cells/well in 1,000 μl of DMEM with 0.4% FBS, and allowed to adhere for 30min at 37°C. Nonadherent cells were removed by gentle washing twice with PBS, and attached cells were fixed in 4% PFA for 15 min and then stained with 2.3% crystal violet (Sigma-Aldrich, MO, USA) for 10min. Cells were extensively washed, air dried, and then the cells imaged and counted using ImageJ. All experiments were performed in triplicate, and each experiment was repeated at least three times.

***Analysis of gene expression data***

Gene expression data from three different cohorts of patients with breast cancers (Bos et al., 2009; Minn et al., 2005; Wang et al., 2005) were collected from the National Center for Biotechnology Information (NCBI) Gene Expression Omnibus (GEO) website (Barrett, T. et al. 2005; Edgar et al., 2002). The dataset was prepared by pooling data from GSE2034, GSE2603 and GSE12276 and normalizing them together using Robust Multi-chip Average (RMA) algorithm. Patient survival data were carefully annotated for Kaplan-Meier analysis. To derive optimal cut-off values of gene expression levels, they are ordered from low to high and a rising step function was computed to define a threshold by StepMiner algorithm (Sahoo et al., 2007). Gene expression values were converted to high and low levels based on the StepMiner threshold. A noise margin of +/- 0.5 was used around the StepMiner threshold to provide relaxed estimates of the high/low values. A noise margin of +/- 0.5 around StepMiner threshold was used to soften or harden the actual threshold. Time-dependent survival probabilities are estimated with the Kaplan-Meier method (Kaplan EL, 1958) and compared using the log-rank test (Peto et al., 1977). Cox proportional-hazards regression models and life-tables (Cox, 1972) were used to test the statistical interaction between two different genes based on their association with survival outcome. High and low expression patterns for focal genes such as GIV (CCDC88A), K2 (FERMT2) and ITGB1 were compared individually using Kaplan-Meier analyses in R statistical software (R version 3.4.4 2018-03-15). Statistical interaction (synergistic effects) between the focal adhesion genes GIV (CCDC88A), K2 (FERMT2), ITGB1, TLN1 (talin 1), TNS1 (tensin 1), TENC1 (TNS2, tensin 2), TNS3 (tensin 3), TNS4 (tensin 4), PXN (paxillin), and VCL (vinculin) were measured using interaction terms in Cox proportional hazards regression model on the above pooled breast cancer dataset with survival data. For example, coefficient of interaction terms was computed from Cox regression model for CCDC88A and FERMT2 as follows:

$$\mathrm{h(t)} = \mathrm{h_0(t)exp(a1} * \mathrm{CCDC88A} + \mathrm{a2} * \mathrm{FERMT2} + \mathrm{a3} * \mathrm{CCDC88A} * \mathrm{FERMT2})$$

where $\mathrm{h(t)}$ is the hazard rate at time $\mathrm{t}$ for an individual; $\mathrm{a1, a2,}$ and $\mathrm{a3}$ are the regression coefficients; $\mathrm{h_0(t)}$ is the baseline hazard; two indicator variables $\mathrm{CCDC88A}$ and $\mathrm{FERMT2}$ (high expression = 1 else 0) that had only additive or interactive effects; $\mathrm{a3}$ is the coefficient of interaction terms.

**Data analysis and other methods**

All experiments were repeated at least three times, and results were presented either as one representative experiment or as average ± SD. Statistical significance was assessed with the Student's t test. Statistical significance between datasets with three or more experimental groups was determined using one-way analysis of variance (ANOVA) including a Tukey's test for multiple comparisons. *p < 0.05, **p < 0.01, ***p < 0.001, ****p < 0.0001.

### SUPPLEMENTARY FIGURES AND LEGENDS

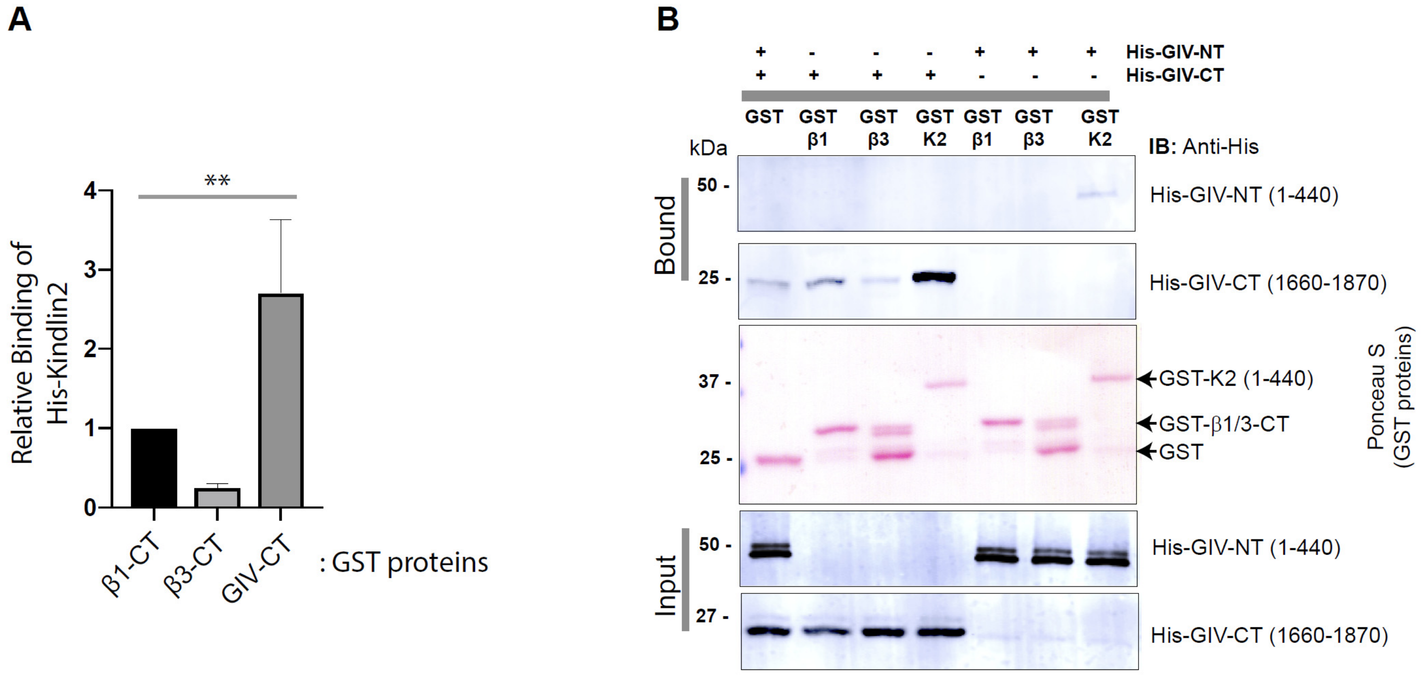


**Figure S1 [Related to Figure 1]**:

**The C-terminus, but not the terminus of GIV binds directly and strongly to Kindlin-2, and to a much weaker degree to β1-integrin tail**.

- **A.** Bar graphs display the relative binding of His-Kindlin 2 to various GST proteins in **Fig 1C**. Error bar = S.E.M (n = 3); **p=<0.01.
- **B.** GST pulldown assays were carried out using His-tagged GIV--CT or NT proteins (~ 3 µg) and GST-tagged β1-integrin tails and Kindlin protein immobilized on glutathione beads. Equal loading of input (lower panels) and bound (upper panels) proteins were visualized by immunoblotting with anti-His antibody. GST proteins were visualized using Ponceau S staining.

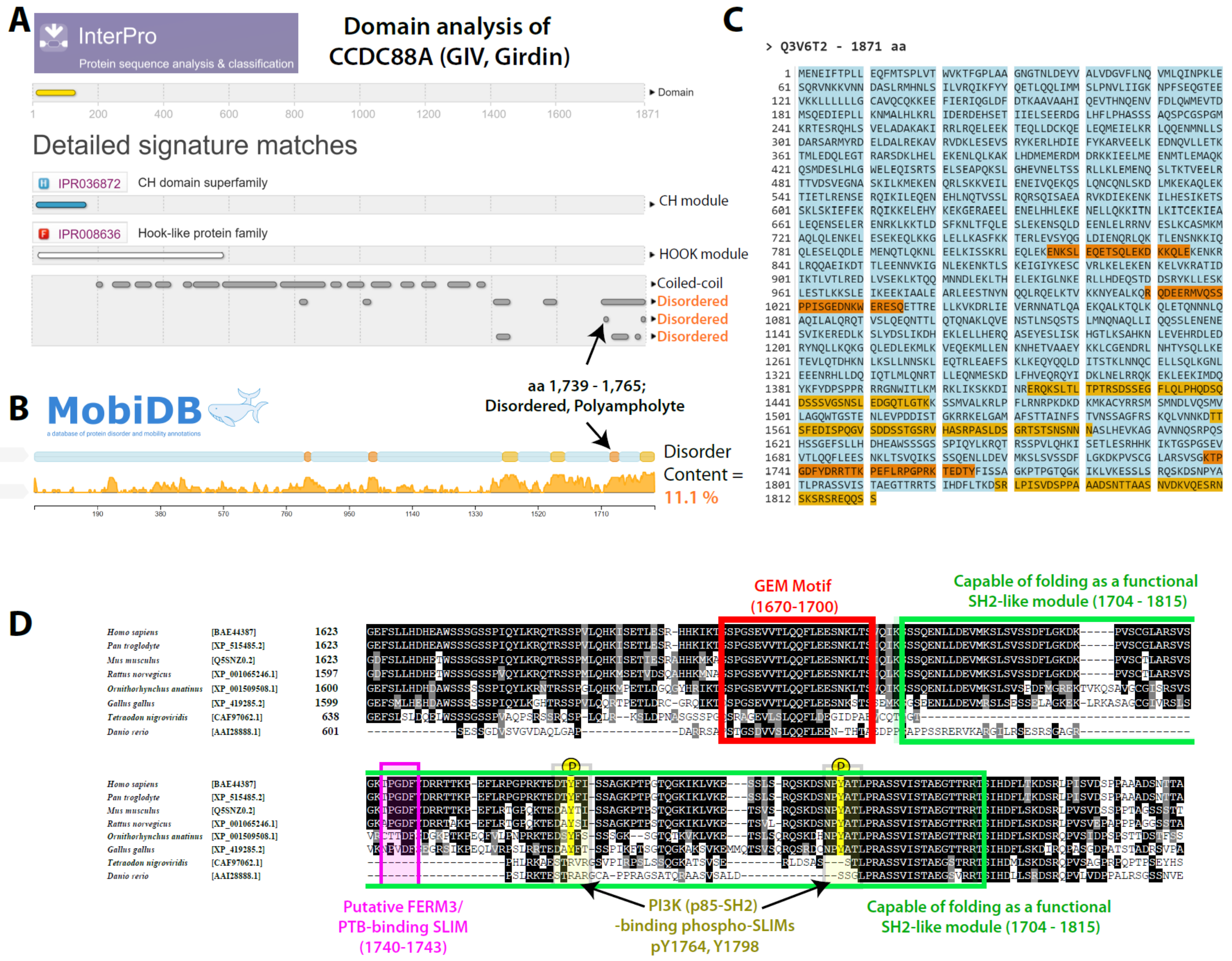


**Figure S2 [Related to Figure 2]**:

**GIV's C-terminal ~ 300-400 amino acid stretch is largely disordered, and aa 1739 – 1765 is predicted to have the highest disorderliness. A.** GIV's sequence was analyzed for structural modularity using InterPro [https://www.ebi.ac.uk/interpro/], a web based program that provides functional analysis of proteins by classifying them into families and predicting domains and important sites by finding matches against a number of member databases into a single searchable resource. Analysis showed the previously known and validated presence of N-terminal coiled coil and Hook domains. The C-terminus is predicted to have disordered segments, which was also confirmed by a secondary web-based applications, MobiDB [http://mobidb.bio.unipd.it/], which represents the disorder content as 2D plots (**B**). The intensity of orange color denotes the degree of disorderliness. **C**. Sequence of GIV with highlighted stretches of disorderliness shows highest disorderliness in the stretch aa 1739-1765, within which the putative FERM3-PTB binding sequence PGxF is located. **D**. Sequence alignment of GIV's C-terminus showing a complete catalog of previously validated short linear motifs (SLIMS) and modules; the position of the newly identified FERM3-PTB binding SLIM (aa 1740-1743) is shown in pink. Because it is located within GIV's SH2-like module, it is unlikely that GIV would bind RTKs and Kindlin (or other PTB modules) simultaneously. Because multiple phosphosites flank this SLIM, it is likely that binding of GIV to PTB modules are regulated by post-translational modifications.

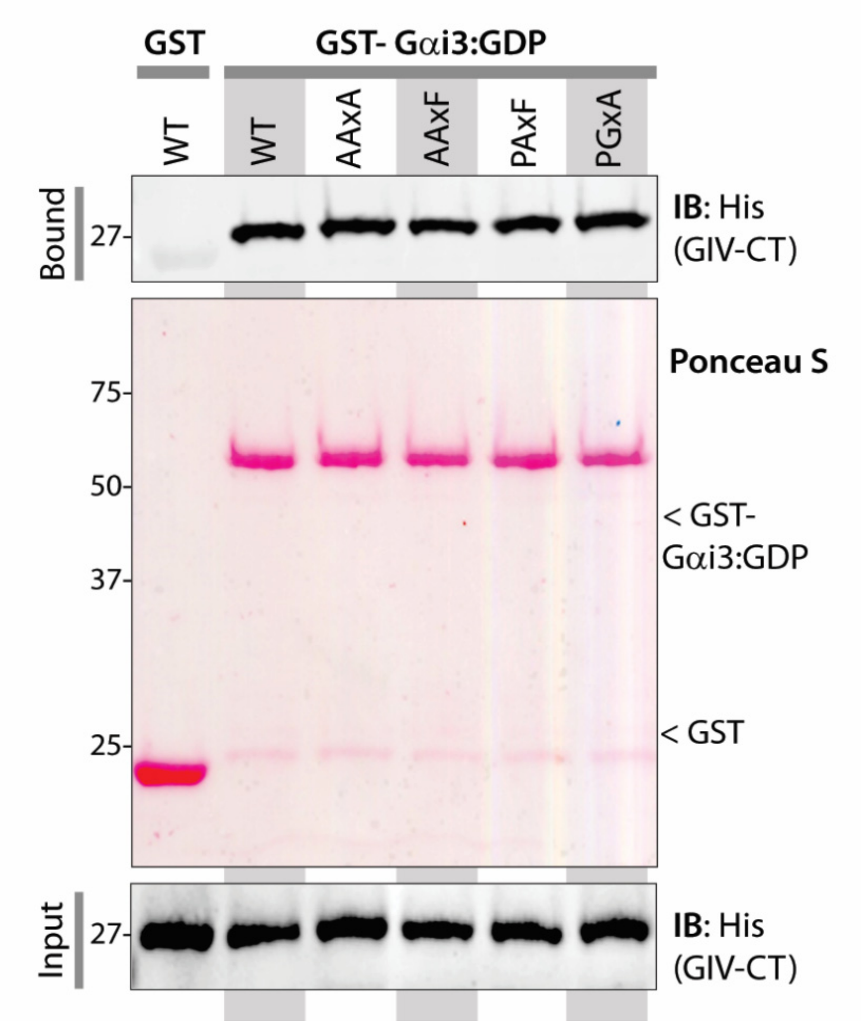


**Figure S3 [Related to Figure 2]**:

**Mutations that impair GIV•K2 interaction do not perturb the GIV•Gαi interaction.** GST pulldown assays were performed using of His-GIV-CT WT or various mutants targeting its PGxF sequence and GST-Gαi3 (pre-loaded with GDP). Equal loading of GST proteins was confirmed by Ponceau S staining.

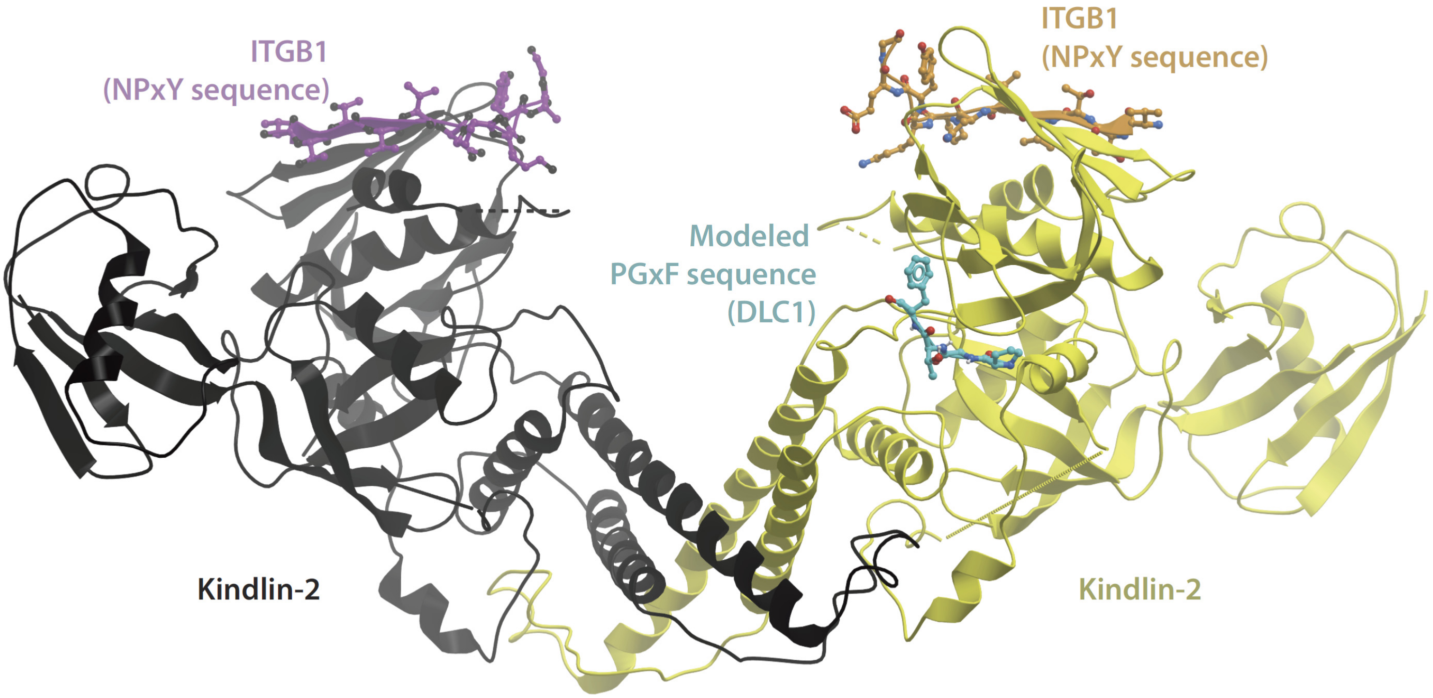


**Figure S4 [Related to Figure 2]**:

**The proposed GIV-PGxF sequence binding surface is accessible in dimerized Kindlin-2.** Figure shows the solved structure of K2 dimers (black and yellow) bound to ITGB1 tail (magenta and orange). The proposed interface with PGxF motif (teal blue) is shown on one of the monomeric units (yellow) when engaged as a dimer. This model was created using the solved structures of ITGB1-bound K2 (PDB:5XQ0) and DLC1(PGxF)•tensin complex structure (PDB:2LOZ).

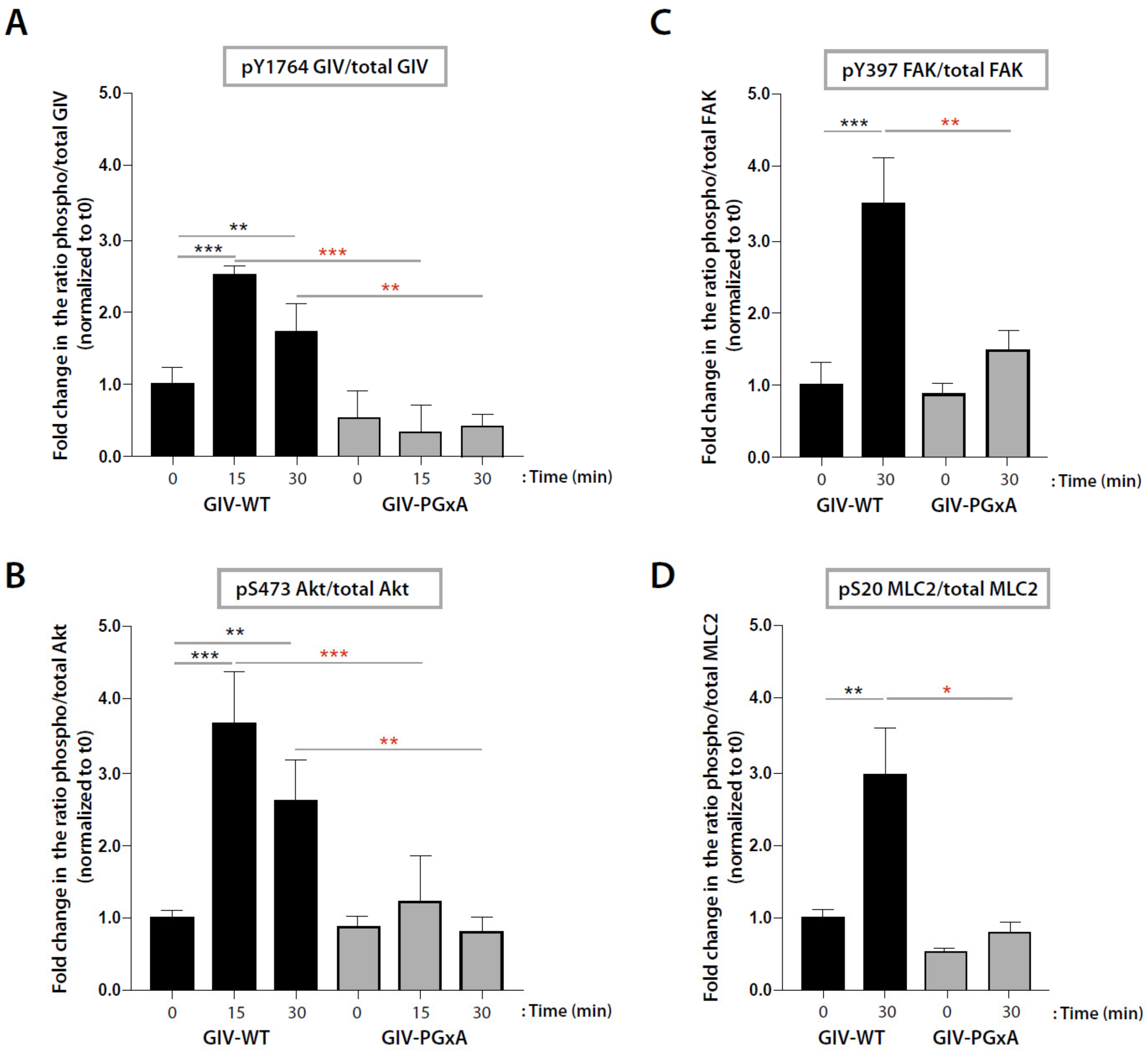


**Figure S5 [Related to Figure 4]**:

**Mutations that impair GIV•K2 interaction suppress major signaling pathways previously shown to be triggered downstream of Integrin β1.** Bar graphs display quantification of 3-4 independent experiments measuring the ratio of phospho/total GIV (**A**), Akt (**B**), FAK (**C**) and MLC2 (**D**). Error bars = S.E.M. *$p < 0.05$, **$p < 0.01$, ***$p < 0.001$. See **Fig 4J-L** for representative blots.
